# Hypoxia-induced histone methylation and NF-κB activation in pancreas cancer fibroblasts promote EMT-supportive growth factor secretion

**DOI:** 10.1101/2025.01.30.635486

**Authors:** Karl M. Kowalewski, Sara J. Adair, Anne Talkington, Jason J. Wieder, Jason R. Pitarresi, Kia Perez-Vale, Britney Chu, Sepideh Dolatshahi, Rosalie Sears, Ben Z. Stanger, Todd W. Bauer, Matthew J. Lazzara

## Abstract

The pancreatic ductal adenocarcinoma (PDAC) tumor microenvironment contains hypoxic tissue subdomains and cancer-associated fibroblasts (CAFs) of multiple subtypes that play tumor-promoting and -restraining roles. Here, we demonstrate that hypoxia promotes an inflammatory-like CAF phenotype and that hypoxic CAFs selectively promote epithelial-mesenchymal transition (EMT) in PDAC cancer cells through growth factor-mediated cell crosstalk. By analyzing patient tumor single-cell transcriptomics and conducting an inhibitor screen, we identified IGF-2 and HGF as specific EMT-inducing growth factors produced by hypoxic CAFs. We further found that reactive oxygen species-activated NF-κB cooperates with hypoxia-dependent histone methylation to promote IGF-2 and HGF expression in hypoxic CAFs. In lineage-traced autochthonous PDAC mouse tumors, hypoxic CAFs resided preferentially near hypoxic, mesenchymal cancer cells. However, in subcutaneous tumors engineered with hypoxia fate-mapped CAFs, once-hypoxic re-oxygenated CAFs lacked a spatial correlation with mesenchymal cancer cells. Thus, hypoxia promotes reversible CAF-malignant cell interactions that drive EMT through druggable signaling pathways.

**One-sentence summary:** We show that hypoxic fibroblasts in pancreas cancer leverage histone methylation and ROS-mediated NF-κB activation to produce growth factors that drive epithelial-mesenchymal transition in malignant cells, demonstrating how tumor stromal features cooperate to initiate a signaling process for disease progression.

## INTRODUCTION

The pancreatic ductal adenocarcinoma (PDAC) tumor microenvironment (TME) is notoriously fibrotic and hypoxic, with a dense stroma that is poorly vascularized and crowded with cancer-associated fibroblasts (CAFs) that account for up to 80% of tumor cellularity (*1, 2*). PDAC tumor enrichment for hypoxic transcriptomic-proteomic signatures or for the presence of CAFs correlates with reduced patient survival (*3–6*), and several mechanisms have been proposed for each effect. Hypoxia promotes PDAC chemoresistance through elevated glutaminolysis, tumor invasiveness by driving glycolytic metabolism, and immunosuppression by recruiting myeloid-derived suppressor cells (*7–9*). CAFs promote PDAC progression by secreting exosomes that support cancer cell proliferation, impeding natural-killer cell function via IL-6 secretion, and promoting chemoresistance by producing growth factors (*10–12*). CAF subpopulations that restrain tumor growth have also been identified, such as those that express meflin and impede collagen matrix remodeling (*13, 14*). Hypoxia and CAFs may also affect PDAC progression through a potentially shared ability to drive epithelial-mesenchymal transition (EMT), which is associated with poor tumor differentiation, chemoresistance, metastasis, and reduced patient survival (*4, 15–17*). Hypoxia drives a cell-autonomous EMT in PDAC cancer cells (i.e., malignant ductal cells) (*18–20*), and CAFs can drive EMT by secreting cytokines including HGF and IL-6 (*21–24*). Interestingly, both cancer cells and CAFs occupy hypoxic regions in PDAC tumors (*2, 25*), but the potential for hypoxia to modulate the ability of CAFs to promote cancer cell EMT has not been investigated.

Hypoxia-inducible factors (HIFs) are one of the most well studied cellular mechanisms of oxygen sensing and have functional roles in CAFs. In addition to oxidative regulation, HIF abundance is strongly controlled by several kinase signaling pathways (*26, 27*), making the regulation of HIF and its downstream effects complex. In PDAC CAFs, HIF2α mediates paracrine signaling that causes M2 macrophage polarization and resultant immune suppression (*28*). In breast cancer, HIF1α promotes a glycolytic CAF phenotype that supports cancer cell protein biosynthesis (*29*). Oxygen sensing can also arise through epigenetic mechanisms. Indeed, low oxygen tension impairs the activities of histone demethylases KDM2A, KDM5A, and KDM6A (*19, 30, 31*). The resultant increases in histone methylation marks, including those on histone H3 lysine 36, histone H3 lysine 4, and histone H3 lysine 27, strongly regulate gene expression by increasing or decreasing promoter accessibility (*30, 31*). For example, histone H3 lysine 27 trimethylation (H3K27me3) inhibits WNT5A and IGF-2 expression in gastric cancer CAFs (*32*) and reduces CAF cytokine production in high-grade serous ovarian carcinoma (*33*). H3K36me2 promotes IGF-2 and TGFα expression in multiple myeloma (*34*). Whether HIFs or impaired histone demethylase activities impact PDAC CAF responses to hypoxia, especially in ways that alter ductal cell EMT, is unknown.

A potentially related issue is the degree to which hypoxia promotes the switching of CAFs among subtypes that arise from pancreatic stellate cells (PSCs) and other mesenchymal lineages (*35, 36*). Myofibroblastic CAFs (myCAFs) arise in response to cancer cell-initiated TGFβ/SMAD (*36, 37*) and support immune response through type-1 collagen expression by reducing myeloid-derived suppressor cell infiltration (*38*). Inflammatory CAFs (iCAFs) arise in response to cancer cell-initiated IL1α/JAK/STAT signaling (*37*) and facilitate PDAC growth and invasiveness through paracrine signaling interactions that activate the JAK/STAT pathway in neoplastic cells (*24, 39*). Both subtypes have the potential to drive EMT in ductal cells. myCAFs remodel the PDAC extracellular matrix (ECM) (*36, 38*) and may regulate EMT through ECM stiffening and collagen fiber alignment (*35, 40*). Compared to myCAFs, iCAFs are generally more secretory and express cytokines including IL-6 (*41*), which can induce ductal cell EMT in PDAC (*22, 42*). Interestingly, hypoxia promotes expression of the iCAF subtype-inducing cytokine IL1α in murine-derived PDAC organoids (*43*), and HIF1α expression promotes the iCAF phenotype in murine PSCs (*44*). Additional research is needed to test these findings using human PDAC CAFs. Two additional hypoxia-responsive biochemical switches, NF-κB activation (*45, 46*) and histone modifications (*19, 30, 31*), remain unexplored in the regulation of PDAC CAF heterogeneity.

Here, we demonstrate that hypoxic CAFs execute a reversible phenotype switch in response to cooperating transcriptional and epigenetic effects and that the resultant iCAF-like cells potently drive EMT by activating cancer cell receptor tyrosine kinases. In patient tumors, iCAFs were enriched for hypoxia-regulated transcripts, and specific EMT-promoting ligand-receptor interactions were enriched between iCAFs and cancer cells. These findings are relevant in mouse models of PDAC, where hypoxic cancer cells proximal to hypoxic CAFs displayed increased mesenchymal characteristics compared to those distal to hypoxic iCAFs. The mechanisms nominate several ways for controlling PDAC TME-mediated interactions to control chemoresistance engendered by EMT.

## RESULTS

### iCAFs promote EMT in PDAC cancer cells

To predict paracrine signaling interactions between iCAFs or myCAFs and PDAC cancer cells, we analyzed previously acquired single-cell RNA sequencing (scRNA-seq) data from six PDAC patients (*41*). As described in *Methods,* we aggregated three distinct ductal cell clusters identified by Elyada et al. to model the cancer cell population. CAFs were hierarchically clustered based on expression of 452 growth factors and cytokines compiled by the Molecular Signatures Database (*47*), with only those cytokines expressed by ≥ 10 cells retained for analysis (102 genes retained). The heatmap in **Figure 1A** shows that distinct secretome profiles were found for iCAFs and myCAFs that may relate to their function in tumors. For example, PDAC iCAFs expressed IL-6 and CXCL12, which promote cancer cell invasion and metastasis (*48, 49*). PDAC myCAFs expressed MDK and MIF, which promote PDAC cell proliferation (*50, 51*). To predict ligand-receptor interactions that may occur between iCAFs or myCAFs and ductal cells, we utilized CellPhoneDB (*52*) (**Figure 1B**) and CellChat (*53*) (**Supp Fig S1A**). CellPhoneDB results were filtered for the top 50 CAF cell (sender) to ductal cell (receiver) interactions. CellPhoneDB indicated that iCAFs communicate with ductal cells via ligands including FGF7/9, IGF1/2, IL-6, and CXCL12 while myCAFs utilize factors such as PDGFD and MDK. CellChat analysis corroborated that iCAFs produce IGFs, FGFs, and CXCL cytokines and further identified that myCAFs produce periostin and MIF, which were not found by CellPhoneDB analysis (**Supp Fig S1A**). Using the NicheNet database (*54*), we computed predicted regulatory potentials of CellPhoneDB-nominated ligands by generating an integrated score for mesenchymal genes in a pan-cancer EMT ontology derived from patient samples (*55, 56*). This was done for the 21 transcripts shared between the CellPhoneDB analysis and the NicheNet database (**Figure 1C**). iCAF-secreted factors, including IGF1/2, AXL ligands (PROS1 and GAS6), and EGFR ligands (AREG and HBEGF), tended to have higher predicted EMT-regulatory potentials than myCAF-secreted ligands, which led us to hypothesize that iCAFs are more likely to promote EMT than myCAFs. The diversity of growth factors identified is consistent with the known involvement of multiple, independent pathways in EMT regulation (*57*).

**Figure 1.**
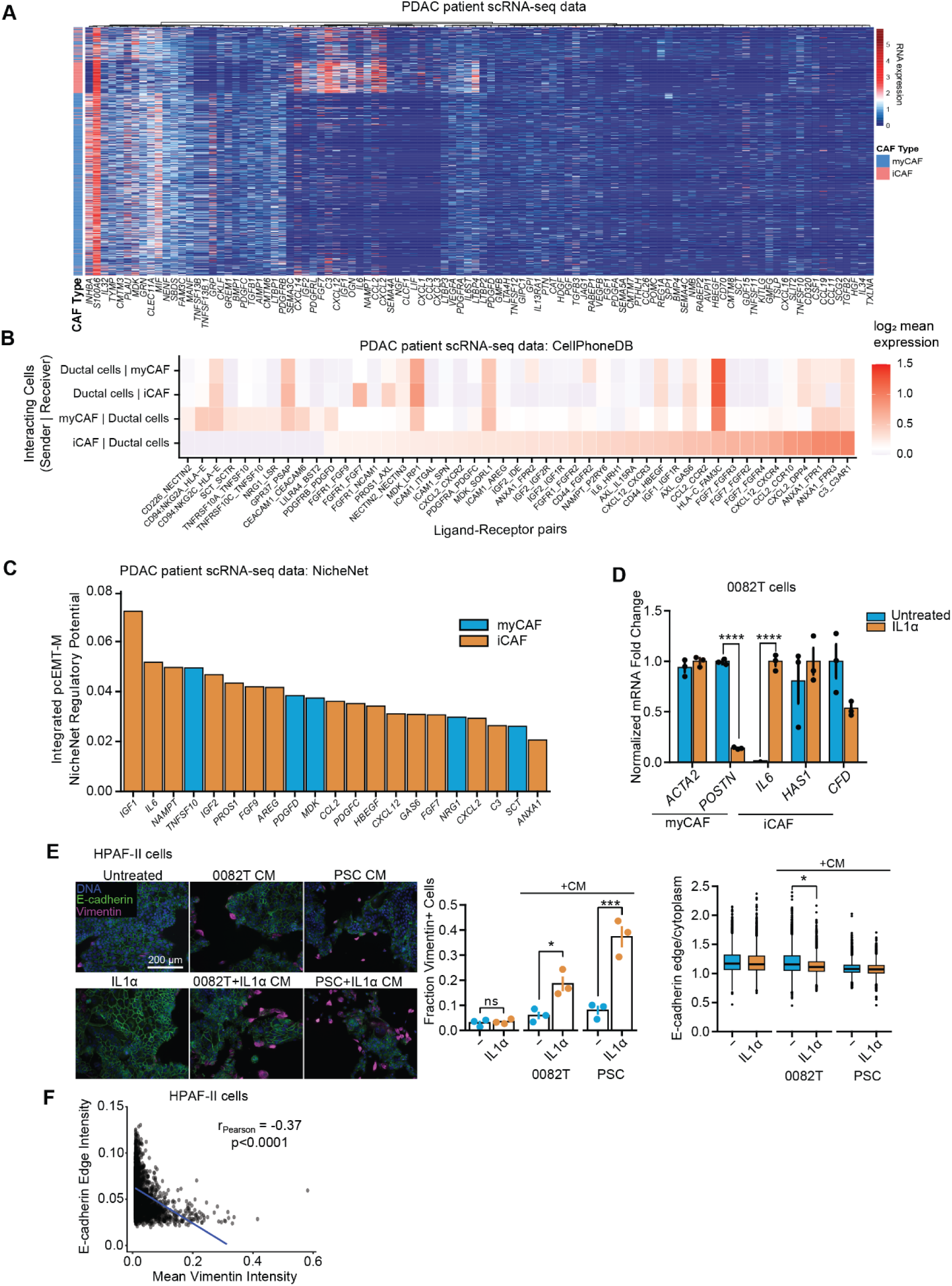
Pancreas cancer iCAFs are enriched for expression of EMT-inducing cytokines. **A)** Using annotated scRNA-seq data from PDAC patient tumors (*41*), PDAC CAFs were hierarchically clustered using Ward’s method based on expression of growth factors and cytokines. **B)** Using the same data analyzed in panel (A), ligand-receptor interactions between iCAFs/myCAFs and cancer cells were predicted using CellPhoneDB. **C)** Growth factor and cytokine regulatory potential for mesenchymal genes in the pan-cancer EMT (pcEMT-M) signature was determined for ligands nominated by CellPhoneDB analysis. Integrated scores for each ligand were calculated by summing the regulatory potential from the NicheNet database for each gene contained in the pcEMT-M gene set. **D)** 0082T CAFs were cultured for 120 hr ± IL1α (10 ng/mL), and qRT-PCR was performed on extracted RNA for the indicated myCAF or iCAF markers. *GAPDH* was used for normalization. n=3, t-test performed per transcript. **E)** HPAF-II cells were treated for 120 hr with IL1α (10 ng/mL) or with conditioned medium (CM) from 0082T cells or PSCs that had been pretreated for 72 hr with or without IL1α (10 ng/mL). Immunofluorescence was then performed on HPAF-II cells for the indicated proteins, and image analysis was performed. One-way ANOVA with Tukey’s multiple comparison test comparing all conditions was performed for vimentin positivity and E-cadherin. **F)** Vimentin and E-cadherin expression were compared for individual cells from the +PSC+IL1α CM condition, and a fit was performed by linear regression. Linear regression t-test. Data are presented as mean ± SEM for bar plots depicting data from 0082T and HPAF-II cells (D, E) or as box-and-whisker plots that represent the first quartile (lower box bound), median (center bound), or third quartile (upper box bound) (E-cadherin measurements in panel E). * *p* <0.05, ** *p* < 0.01, *** *p* < 0.001, **** *p* < 0.0001

To test the hypothesis that iCAFs are more potent drivers of PDAC cancer cell EMT than their more myofibroblastic counterparts, we used IL1α to drive a transition toward the iCAF phenotype in human PSCs and the 0082T immortalized human PDAC CAF line. To confirm an IL1α-induced phenotypic transition, we tested a subset of subtype markers by qRT-PCR. IL1α reduced expression of the myCAF marker *POSTN* and increased expression of *IL6* in 0082T CAFs, while other iCAF (*HAS1* and *CFD*) and myCAF (*ACTA2*) markers were unaffected, suggesting that additional factors may be necessary to achieve a more complete iCAF transition (**Figure 1D**). The iCAF and myCAF phenotypes were characterized from a heterogeneous CAF population by scRNA-seq data (*41*), so a single cell line may not capture all features of the iCAF signature. Therefore, we refer to this phenotype as “iCAF-like.” At the protein level, expression of the iCAF marker IL-6 was enhanced in 0082T CAFs and human PSCs treated with IL1α (**Supp Fig S1B**). To test the ability of iCAF-like cells to promote EMT, conditioned medium (CM) from 0082T cells or PSCs with or without IL1α pretreatment was used to treat HPAF-II human PDAC cancer cells. Compared to no pretreatment, IL1α pretreatment of 0082T cells or PSCs led to a 2.6- or 3.8-fold increase, respectively, in HPAF-II expression of the mesenchymal marker vimentin (**Figure 1E**). IL1α treatment did not increase HPAF-II cell vimentin expression, demonstrating that the CM effects were not due to IL1α carryover. In HPAF-II cells treated with PSC+IL1α CM, increased vimentin expression was accompanied by a loss of membranous E-cadherin, another anticipated effect of EMT (**Figure 1F**). Because its changes are more easily quantified, vimentin was used as the primary EMT marker in this study. To test the generality of these findings, we used two primary human CAF lines that exhibited baseline iCAF- or myCAF-like properties, based on qRT-PCR of subtype marker genes (**Supp Figure S1C**). CM from the more iCAF-like CAFs produced a 6.6-fold higher fraction of vimentin-positive HPAF-II cells than did the myCAF-like CAFs (**Supp Figure S1D**).

We next hypothesized that kinase-dependent signaling in cells treated with CAF CM was responsible for the observed EMT, based in part on the established role of MAPK signaling in pancreas and lung cancer EMT (*19, 58, 59*). In lysates of HPAF-II cells treated with PSC+IL1α CM, a phospho-protein array identified increased phosphorylation of ERK1/2, RSK1/2 (ERK substrates), and STAT3 (**Supp Figure S2A**). MEK inhibition effectively reduced HPAF-II cell EMT in response to CM from 0082T cells treated with or without IL1α, confirming a role for ERK in iCAF-induced EMT (**Supp Figure S2B**). Although STAT3 phosphorylation increased in HPAF-II treated with PSC+IL1α CM, STAT3 inhibition failed to antagonize EMT (**Supp Figure S2C**).

### Hypoxia supports CAF-mediated EMT in PDAC cancer cells

Hypoxia is a pervasive feature of PDAC tumors that promotes the iCAF phenotype in murine PSCs (*43, 44*). To test for a connection between hypoxia and the iCAF phenotype in the data from Elyada et al. (*41*) that were analyzed in Figure 1, we performed gene set variation analysis (GSVA) on the CAF cell population. PDAC iCAFs were enriched in Hallmark Hypoxia genes (**Figure 2A**), consistent with conclusions reached for other datasets (*43*). To test the connection between hypoxia and the iCAF phenotype experimentally, 0082T CAFs were cultured in 21% or 1% O_2_ and assessed for CAF subtype marker expression by qRT-PCR. Hypoxia reduced expression of the myCAF marker *POSTN* and increased expression of the iCAF marker *HAS1* (**Figure 2B**). Hypoxia led to the same suppression of *POSTN* observed in response to IL1α in Figure 1D and an elevated expression of *HAS1* expression that was not seen in response to IL1α. Thus, both hypoxia and IL1α promoted a more iCAF-like phenotype, but there were differences in responsive CAF-subtype markers. Furthermore, even though hypoxia failed to reduce expression of *ACTA2* (encodes αSMA), protein expression of the myCAF marker αSMA decreased in CAFs cultured in 1% O_2_, aligning with prior characterization of the iCAF phenotype (**Figure 2C**). The discrepancy between gene and protein expression may result from αSMA depolymerization in response to hypoxia (*60*).

**Figure 2.**
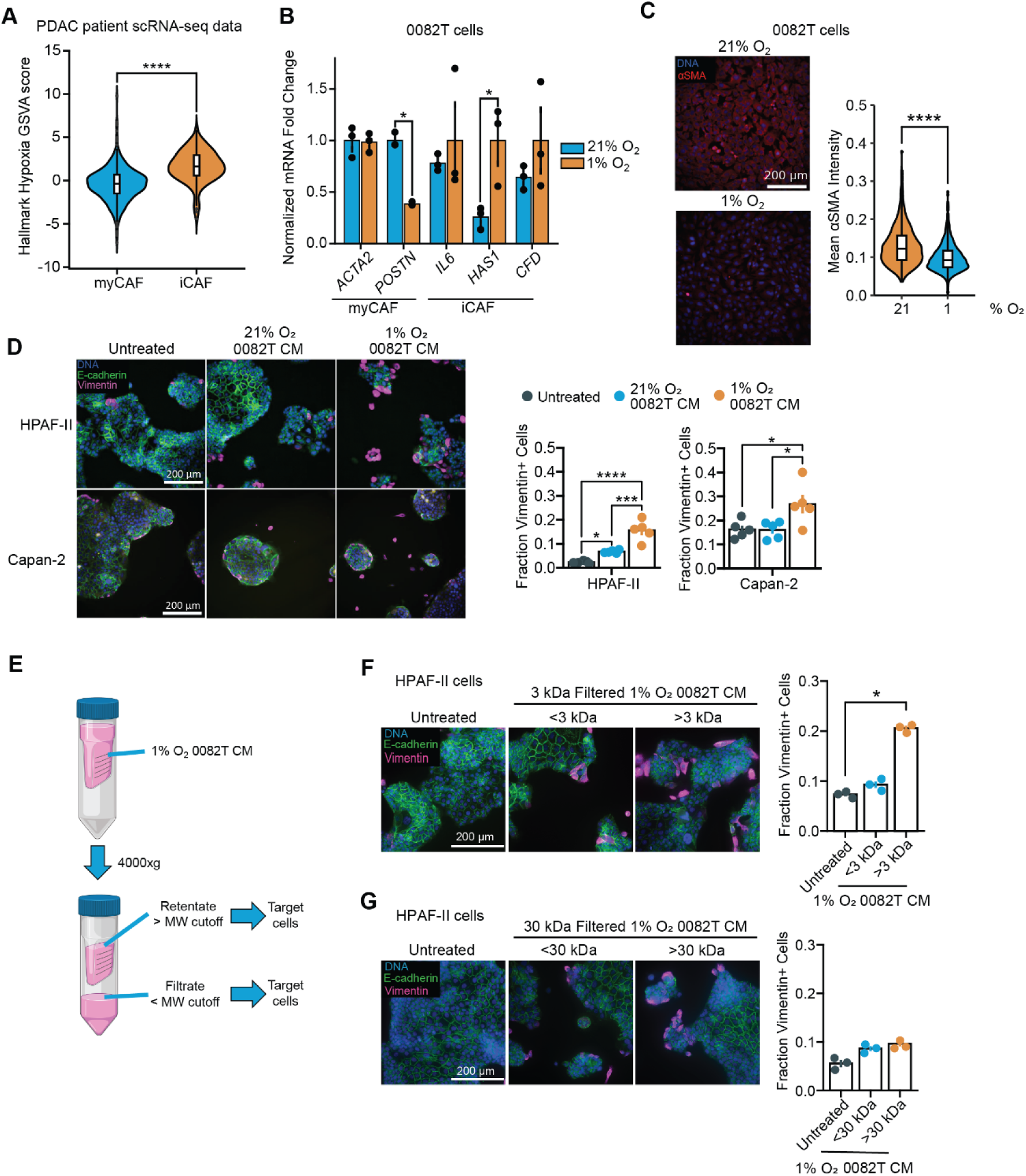
Hypoxia promotes the iCAF phenotype and enhances the ability of CAFs to promote EMT through secreted factors. **A)** Using annotated scRNA-seq data from PDAC patient tumors (*41*), enrichment for Hallmark Hypoxia genes was compared between myCAFs and iCAFs using gene set variation analysis (GSVA). Wilcoxon rank-sum test. **B)** 0082T CAFs were cultured in 21% or 1% O_2_ for 120 hr, and qRT-PCR was performed on extracted mRNA for the indicated myCAF or iCAF transcripts. *CASC3* was used for normalization. n=3, t-test performed per transcript. **C)** 0082T cells were cultured in 21% or 1% O_2_ for 72 hr, and immunofluorescence microscopy was performed for αSMA. n=3, t-test. **D)** HPAF-II and Capan-2 cells were treated for 120 hr with conditioned medium (CM) generated from 0082T CAFs that were cultured in 21% or 1% O_2_. HPAF-II and Capan-2 cells were then analyzed by immunofluorescence microscopy for the indicated proteins. n=4, one-way ANOVA with Tukey’s multiple comparison test. **E)** CM from 0082T cells grown at 1% O_2_ was separated into different molecular-weight fractions using ultrafilters. **F)** CM from 0082T cells grown at 1% O_2_ was separated into retentate and filtrate fractions using 3-kDa filters. HPAF-II cells were treated with the CM fractions for 120 hr, and immunofluorescence microscopy was performed for the indicated proteins. n=3, one-way ANOVA with Tukey’s multiple comparison test. **G)** The experiment described in (F) was performed using 30-kDa filters. n=3, one-way ANOVA with Tukey’s multiple comparison test. Data are presented as mean ± SEM for bar plots (B, D, F, G) or as violin plots overlayed with box-and-whisker plots that represent the first quartile (lower box bound), median (center bound), or third quartile (upper box bound) (A, C). * *p* <0.05, ** *p* < 0.01, *** *p* < 0.001, **** *p* < 0.0001

Because hypoxia promoted an iCAF-like phenotype, we tested if hypoxia also enhanced the ability of CAFs to promote EMT in PDAC cancer cells. 0082T cells were cultured in 21% or 1% O_2_ for 72 hr prior to 48 hr media conditioning at the same oxygen concentrations. As hypothesized, CM from hypoxic 0082T cells roughly doubled the fraction of vimentin-positive HPAF-II cells and caused a 1.5-fold increase in vimentin-positive Capan-2 cells compared to CM from normoxic 0082T cells (**Figure 2D**). Similarly, CM from hypoxic PSCs almost doubled the fraction of vimentin-positive HPAF-II compared to CM from normoxic PSCs (**Supp Figure S3A**).

Hypoxia drives a cell-autonomous EMT in PDAC cell lines, including HPAF-II (*19*), leading to a question of whether hypoxic CAF-cancer cell crosstalk appreciably augments EMT for hypoxic neoplastic cells. We thus tested the effect of CM from hypoxic CAFs on hypoxic cancer cells. Whereas hypoxia alone prompted 12% of HPAF-II cells to become vimentin-positive, the addition of hypoxic CAF CM to the hypoxic culture increased vimentin positivity to 30% (**Supp Figure S3B**). The addition of normoxic CAF CM provided a substantially more modest supplementary effect. These results suggest that secreted factors from hypoxic CAFs cooperated with the cell-autonomous hypoxia-induced EMT. Because hypoxic PDAC cancer cells produce IL1α to promote the iCAF phenotype in murine-derived PDAC organoids (*43*), we hypothesized that coculturing CAFs with PDAC cancer cells in hypoxia would further enhance the ability of hypoxic CAFs to induce EMT. Whereas CM from hypoxic 0082T monocultures promoted EMT in ∼18% of HPAF-II cells, CM from hypoxic cocultures of HPAF-II and 0082T cells promoted EMT in 28% of HPAF-II cells (**Supp Figure S3C**). Thus, CAF-cancer cell crosstalk enhanced the effect of hypoxic CAFs on PDAC cancer cell EMT. CellPhoneDB and NicheNet analysis of ductal cell-secreted ligands suggested that MIF and TNFα produced by hypoxic PDAC cells increase NF-κB target gene expression (**Supp Figure S3D, E**), which may be an influential crosstalk mechanism that supports CAF-mediated EMT.

To test the hypothesis that iCAF-like cells produce soluble factors responsible for the cancer cell EMT that ensues in response to CM, we utilized ultrafilters to fractionate CM (**Figure 2E**). With 3-kDa filters, the retentate (proteins larger than 3 kDa) was primarily responsible for EMT in HPAF-II cells (**Figure 2F**). Most growth factors are larger than 3 kDa, suggesting that CAFs primarily secrete ligands that promote EMT, rather than inducing EMT through nutrient depletion or changes to media pH. With 30-kDa filters, neither the retentate nor filtrate significantly promoted EMT (**Figure 2G**). We interpreted this result to indicate that hypoxic CAF CM contained growth factors smaller and larger than 30-kDa that cooperated to drive the more complete EMT observed with the 3-kDa retentate. The data presented next support this hypothesis.

### Hypoxic CAFs promote cancer cell EMT via IGF-2 and HGF

Because hypoxia increased the ability of CAFs to promote cancer cell EMT, we analyzed transcripts from 0082T cells (cultured in 21% or 1% O_2_) encoding the iCAF-enriched ligands with the highest predicted mesenchymal regulatory scores in Figure 1C. Hypoxia significantly increased expression of *IGF2* and *VEGFA*, with the latter serving as a positive control for hypoxia response (**Figure 3A**). We focused on *IGF2* because its expression was significantly elevated in hypoxic CAFs. Using the scRNA-seq dataset analyzed in Figure 1, we performed consensus clustering based on UCell enrichment scores for Hallmark Hypoxia genes (**Supp Figure S4A, B**). Hallmark Hypoxia-high CAFs displayed higher expression of *IGF2* (and *IGF1*) compared to their less hypoxic counterparts (**Supp Figure S4C**). This inference was validated experimentally by immunofluorescence in 0082T cells, where IGF-2 protein was more abundantly expressed by cells cultured in 1% O_2_ than 21% O_2_ (**Figure 3B**). ELISA demonstrated that an above-background level of IGF-2 could only be detected in one sample collected from hypoxic 0082T cells (**Supp Figure S4D**); we attribute the lack of a consistent ability to detect secreted IGF-2 to the fact that only ∼20% of 0082T cells expressed the growth factor (Figure 3B). In further support of the involvement of IGF-2, we found that the IGF1R inhibitor linsitinib antagonized, but did not completely eliminate, the ability of CM from hypoxic 0082T CAFs to promote EMT in HPAF-II cells (**Supp Figure S4E**). The molecular weight of IGF-2 is 7.5 kDa. Based on our interpretation of the 30-kDa filtration experiment (Figure 2G), at least one growth factor larger than 30-kDa remained unidentified. While TGFβ was not nominated by our scRNA-seq analysis as a hypoxia-responsive growth factor driving EMT, we nonetheless investigated its potential role because hypoxia promotes TGFβ expression in other cell lines (*61, 62*). TGFβR inhibition reduced EMT in response to exogenous TGFβ1 but failed to reduce EMT in response to hypoxic CAF CM. Thus, TGFβ is not an effector of hypoxic CAF-induced EMT, at least for the cell lines investigated (**Supp Figure S4F**).

**Figure 3.**
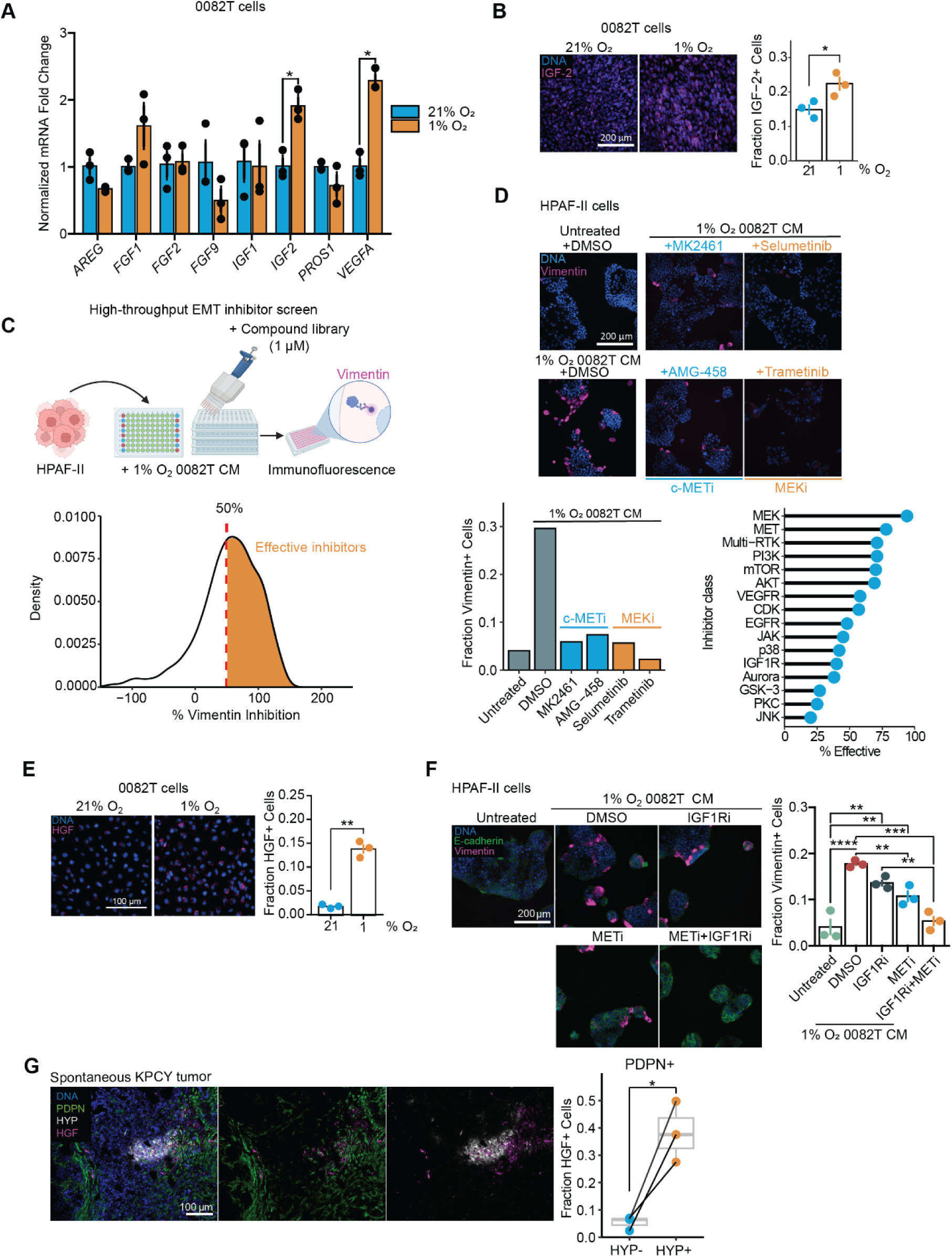
Multiple growth factors and cytokines regulate the ability of hypoxic CAFs to promote EMT in pancreas cancer cells. **A)** 0082T CAF cells were cultured in 21% or 1% O_2_ for 72 hr, and qRT-PCR was performed for the indicated transcripts and normalized to *CASC3*. n=3, t-test. **B)** 0082T CAFs were cultured in 21% or 1% O_2_ for 120 hr, and immunofluorescence microscopy was performed for IGF-2. n=3, t-test. **C)** 430 inhibitors were screened for their ability to counteract hypoxic CAF-induced EMT in HPAF-II cells. HPAF-II cells were treated with hypoxic 0082T CM and inhibitors (1 μM for all compounds) for 120 hr. The ability of each inhibitor to reduce vimentin expression was calculated using positive and negative controls contained on each plate (see Methods). Data are displayed as a plot of the kernel density function (displayed as density) for the percent vimentin inhibition. Inhibitors with >50% reduction in vimentin-positive cells were designated as effective (orange region of the density plot of the inhibitor screen). **D)** Fraction of vimentin-positive HPAF-II cells treated with CM and MET inhibitors (blue) or MEK inhibitors (orange) compared to untreated and CM-only controls. The percentage of effective inhibitors was calculated for each inhibitor class in the inhibitor library. Representative images are shown of effective (i.e., >50% vimentin reduction) MEKi- or METi-treated samples and controls. **E)** 0082T cells were cultured in 21% or 1% O_2_ for 120 hr and assessed for HGF expression by immunofluorescence microscopy. n=3, t-test. **F)** Immunofluorescence microscopy was performed on HPAF-II cells treated with hypoxic 0082T CM with linsitinib (IGF1Ri, 2.5 μM), PHA665752 (METi, 2.5 μM), both, or DMSO. **G)** Immunofluorescence microscopy was performed on KPCY tumor samples to determine fraction of HGF-positive CAFs (PDPN+) in oxygenated (HYP-) or hypoxic (HYP+) tumor regions. n=3, paired t-test. Data are presented as mean ± SEM for bar plots (A, B, E, F) or as a box-and- whisker plot that represent the first quartile (lower box bound), median (center bound), or third quartile (upper box bound) (G). * *p* <0.05, ** *p* < 0.01, *** *p* < 0.001, **** *p* < 0.0001

Given the incomplete effect of linsitinib in Figure S4D, we hypothesized that hypoxic CAFs secrete additional cytokines that drive EMT. To identify candidates, we screened HPAF-II cells treated with hypoxic CAF CM using a library of 430 kinase inhibitors (all at 1 µM) for their ability to antagonize EMT (**Figure 3C**). Inhibitors were designated as effective if they reduced the fraction of vimentin-positive HPAF-II cells by >50% relative to that observed in response to 1% O_2_ 0082T CM (**Figure 3C**). MEK inhibitors, including selumetinib and trametinib, were the most frequently effective at antagonizing EMT, with 16 out of 17 MEK inhibitors in the library reducing vimentin-positive cells by >50% (**Figure 3D**). MET inhibitors, including MK2461 and AMG-458, achieved the second-highest efficacy rate, with 7 out of 9 MET inhibitors in the library effectively reducing the number of vimentin-positive cells (**Figure 3D**). HGF, the ligand for MET, is a well-known and potent EMT driver (*21*), leading us to hypothesize that hypoxia enhances HGF expression in CAFs. 0082T cells cultured in 1% O_2_ displayed significantly higher HGF expression than those cultured in 21% O_2_, as determined by immunofluorescence imaging (**Figure 3E**). Interestingly, reduced oxygen tension in 0082T cells produced no change in *HGF* transcripts (**Supp Figure S5A**), consistent with the absence of *HGF* among the growth factors nominated by the analysis in Figure 1. Hypoxia also reduced the apparent presence of secreted HGF in CM (**Supp Figure S5B**). Given the prioritization of MET by the inhibitor screen and the potency of HGF for EMT, however, we interpret these results to indicate collectively that the export of HGF from CAFs may be forestalled by hypoxia (explaining its intracellular accumulation without increased transcript abundance) but that even the reduced concentrations of secreted HGF are necessary for complete PDAC cell EMT. Moreover, HGF has a molecular weight of 84 kDa, consistent with our hypothesis that CAFs produced an EMT-inducing growth factor larger than 30 kDa. Treating HPAF-II cells with the MET inhibitor PHA665752 reduced EMT in response to CM from hypoxic 0082T cells and exhibited a cooperative effect with the IGF1R inhibitor linsitinib (**Figure 3F**). MET inhibition also effectively reduced EMT in HPAF-II cells treated with hypoxic PSC CM (**Supp Figure S5C**). We additionally confirmed that recombinant HGF and IGF-2 cooperated to promote EMT in HPAF-II (**Supp Figure S5D**).

### CAFs occupy hypoxic regions in PDAC tumors and display enhanced HGF expression

To study the effects of hypoxia on CAFs *in vivo*, we first confirmed the presence of CAFs in hypoxic tumor regions in Kras^LSLG12D^, p53^LSL-R172H^, Pdx1-Cre, Rosa26^LSL-YFP^ (KPCY) mice using pimonidazole (Hypoxyprobe), a reagent that binds to reduced thiols in tissue proteins at low oxygen tension, and podoplanin, a pan-CAF marker (**Figure 3G**). On average, ∼10% of CAFs were found in hypoxic tumor domains (**Supp Figure S5E**). CAFs that occupied regions of hypoxia in KPCY tumors displayed 3-fold greater HGF positivity than CAFs in oxygenated areas (**Figure 3G**). Similar results were obtained using subcutaneous tumors formed by implanting KPCY-derived cell lines (**Supp Figure S5F**).

### Hypoxia activates NF-κB signaling via reactive oxygen species in CAFs

We next sought to identify the signaling mechanisms leading to the hypoxia-induced iCAF-like phenotype. Because NF-κB supports the iCAF phenotype (*37*) and is a common enhancer of cytokine expression [e.g., (*63*)], we hypothesized that hypoxia activates NF-κB signaling to promote CAF production of HGF and IGF-2. Indeed, in 0082T cells, hypoxia increased the expression (**Figure 4A**) and nuclear accumulation (**Figure 4B**) of the NF-κB p65 subunit, consistent with elevated pathway activity. Pretreating hypoxic CAFs with an IKK inhibitor impaired the ability of CAF CM to drive EMT in HPAF-II cells (**Figure 4C**), demonstrating a functional role for CAF NF-κB activity in cytokine-regulated crosstalk with cancer cells. Similarly, doxycycline-inducible shRNA-mediated knockdown of NF-κB (p65) in 0082T cells impaired the ability of hypoxic 0082T CM to drive EMT in HPAF-II cells (**Supp Figure S6A, B**). Furthermore, IKK inhibition reduced HGF expression in hypoxic CAFs (**Supp Figure S6C**). Thus, hypoxia-induced NF-κB pathway activity in CAFs promotes expression of EMT-driving growth factors.

**Figure 4.**
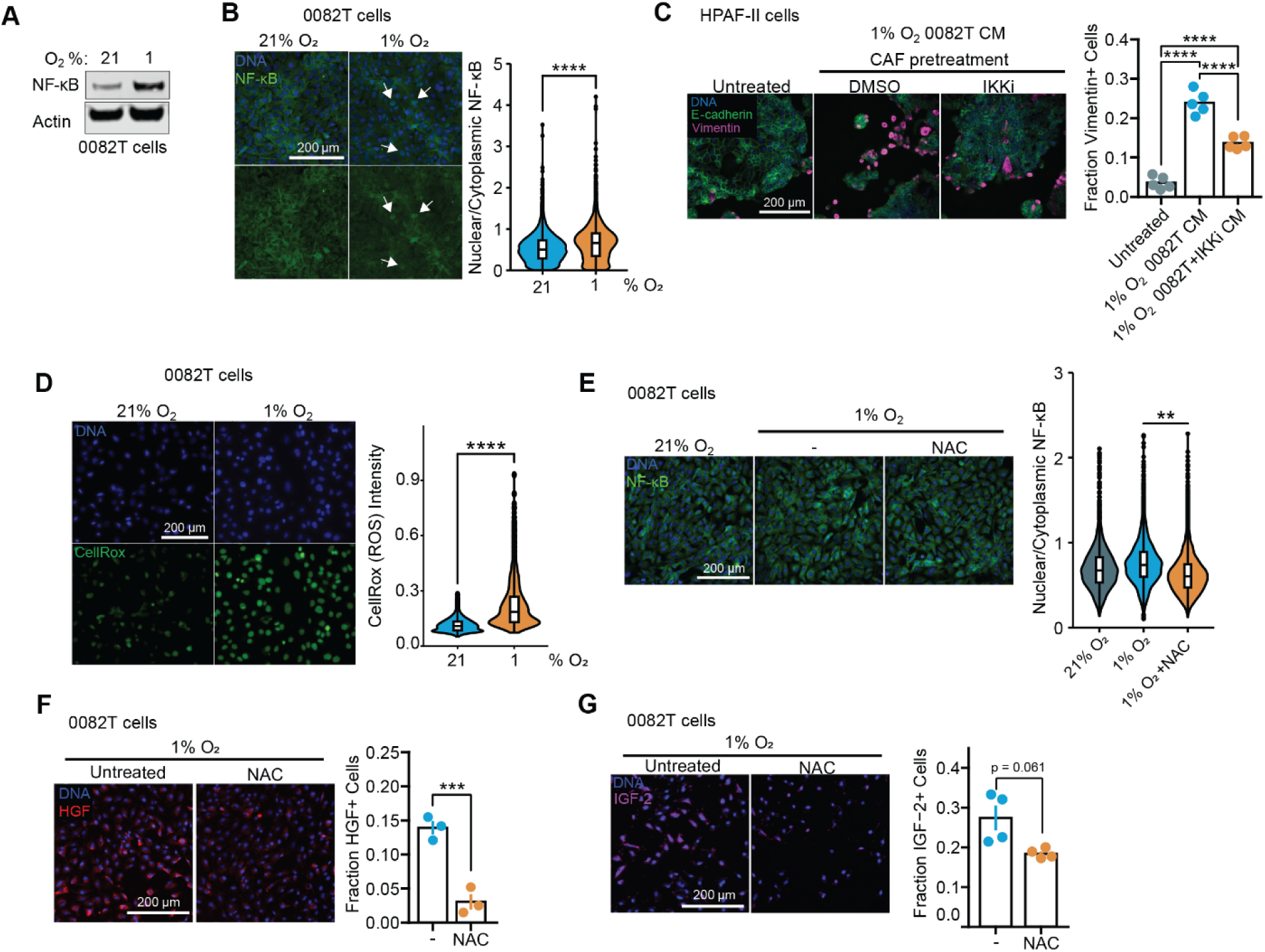
Hypoxia activates NF-κB signaling through reactive oxygen species to promote IGF-2 and HGF expression. **A)** 0082T CAF cells were cultured at 21% or 1% O_2_ for 120 hr, and lysates were subjected to western blotting for the indicated proteins. Blot image representative of n=3. **B)** 0082T cells were cultured in 21% O_2_ or 1% O_2_ for 120 hr, and immunofluorescence microscopy was performed on for NF-κB. n=3, t-test. **C)** HPAF-II cells were treated for 120 hr with CM generated from hypoxic 0082T CAFs pretreated with DMSO or 0.5 μM IKK-16 (IKKi) for 72 hr prior to media conditioning. Immunofluorescence microscopy was performed for the indicated proteins. n=5, one-way ANOVA with Tukey’s multiple comparison test. **D)** 0082T CAFs were cultured in 21% or 1% O_2_ for 120 hr and then analyzed using the CellRox reactive oxygen species (ROS) detection kit. n=3, t-test. **E)** 0082T CAFs were cultured in 21% or 1% O_2_ +/- N-acetyl-cysteine (NAC) for 120 hr. Immunofluorescence microscopy for NF-κB was performed. n=3, ANOVA and Games-Howell post-hoc test. **F)** and **G)** 0082T CAFs were cultured in 1% O_2_ ± NAC for 120 hr. Immunofluorescence microscopy for HGF (**F**) or IGF-2 (**G**) was performed. n=3, t-test. Data are presented as mean ± SEM for bar plots (C, F, G) or as violin plots overlayed with box-and-whisker plots that represent the first quartile (lower box bound), median (center bound), or third quartile (upper box bound) (B, D, E). * *p* <0.05, ** *p* < 0.01, *** *p* < 0.001, **** *p* < 0.0001

Hypoxia causes the accumulation of reactive oxygen species (ROS) (*64*), which can disinhibit NF-κB through effects on the NF-κB inhibitor IκB (69) or IKK (70). Accordingly, we probed for hypoxia-induced ROS accumulation using the fluorescent dye CellRox. CAFs exhibited increased ROS abundance after 5 days of hypoxia (**Figure 4D**). Moreover, the ROS scavenger N-acetyl-cysteine (NAC) reduced NF-κB nuclear accumulation (**Figure 4E**) and suppressed HGF and IGF-2 expression (**Figure 4F, G**) in hypoxic CAFs. Thus, hypoxia enhanced CAF expression of HGF and IGF-2 through a ROS-mediated mechanism that requires NF-κB.

Lactate dehydrogenase (LDHA) is a prominent ROS-generating enzyme (*65*), and our analysis of scRNA-seq patient tumor data demonstrated that transcripts encoding LDHA were elevated in hypoxic CAFs (**Supp Figure S7A**). Other enzymes, especially the NADPH oxidases (NOX), also produce ROS (*66*). However, in patient tumors, transcripts for NOX enzymes were either undetected or did not display increased expression in hypoxic CAFs (**Supp Figure S7A**). Consistent with inferences from patient scRNA-seq data, 0082T CAFs cultured in 1% O_2_ displayed higher LDHA expression than those cultured at 21% O_2_ (**Supp Figure S7B**). We thus hypothesized that ROS generation in hypoxic CAFs was mediated by LDHA. CM from hypoxic CAFs treated with an LDHA inhibitor exhibited higher pH than vehicle-treated CAFs based on the color of the phenol red-containing medium, consistent with reduced lactate production (**Supp Figure S7C**). However, LDHA inhibition in hypoxic CAFs failed to impede NF-κB nuclear accumulation (**Supp Figure S7D**) or suppress HGF expression (**Supp Figure S7E**) and did not impact the ability of hypoxic CAF CM to promote EMT in HPAF-II cells (**Supp Figure S7F**). These data suggest that NF-κB activation is independent of LDHA in hypoxic conditions and may instead rely on mechanisms that were not nominated by our analysis of scRNA-seq data.

### Hypoxia influences CAF phenotype through histone methylation

HIF1α regulates IL-6 expression (an iCAF marker) in murine PSCs (*44*), and HIFs are required for oxygen-sensing in a variety of biological scenarios. We thus investigated the role of HIFs in determining hypoxic CAF phenotype using siRNAs, which effectively reduced HIF expression in 0082T CAF cells (**Supp Figure S8A**). HIF1α knockdown did not impair the ability of hypoxic 0082T CM to promote EMT in HPAF-II cells (**Figure 5A**). Neither HIF1α nor HIF2α knockdown reduced HGF or IGF-2 expression in hypoxic CAFs (**Supp Figure S8B, C**). Thus, the CAF hypoxic response leading to expression of EMT-inducing growth factors was HIF-independent.

**Figure 5.**
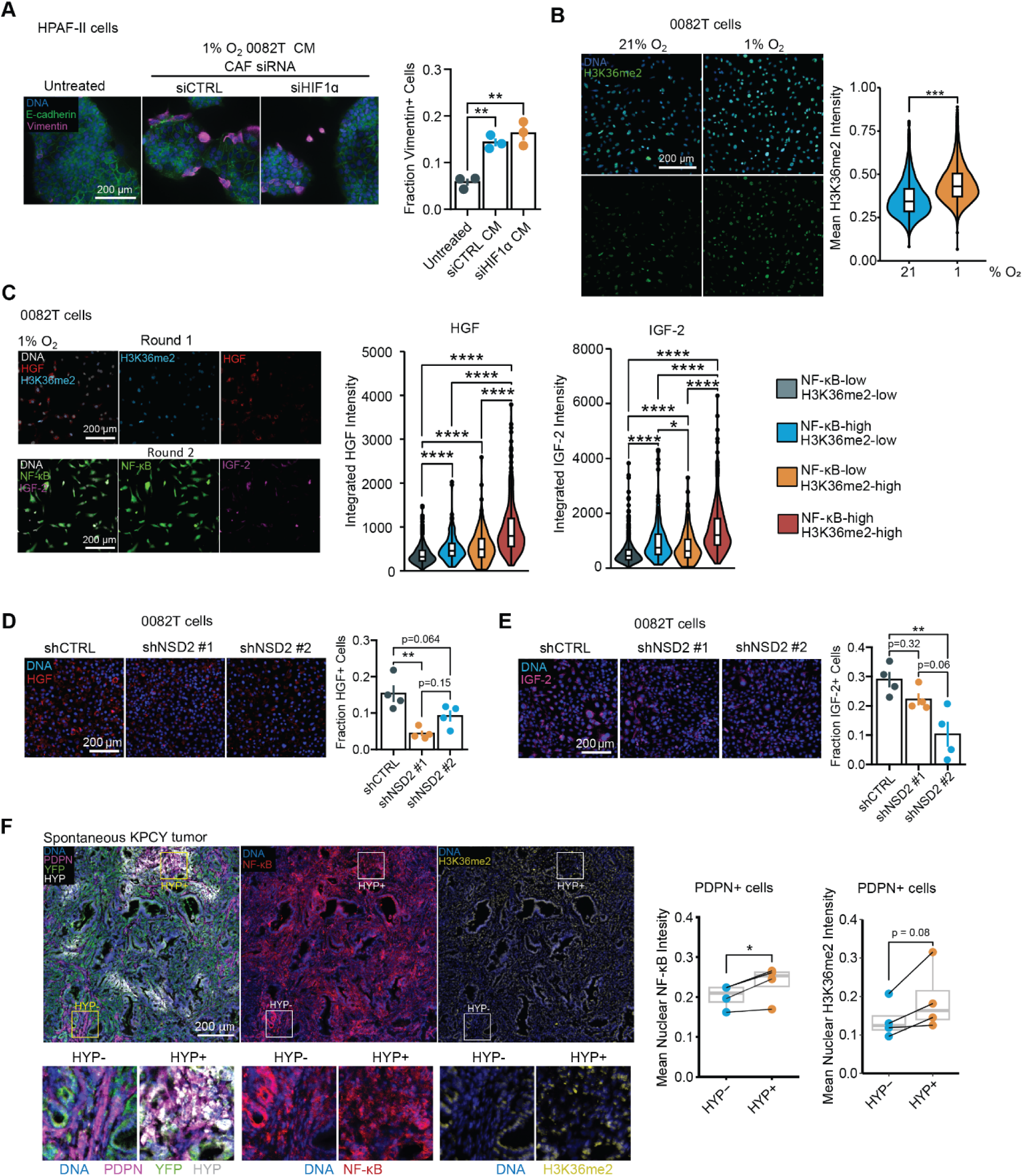
H3K36me2 supports the phenotypic effects of hypoxia on CAFs. **A)** HPAF-II cells were treated for 120 hr with conditioned medium (CM) from 0082T CAFs that were cultured in 1% O_2_ and transfected with control siRNA or *HIF1A*-targeting siRNA. Immunofluorescence microscopy was performed for the indicated proteins. n=3, ANOVA with Tukey multiple comparison test. **B)** 0082T CAFs were cultured in 21% or 1% O_2_ for 120 hr, and immunofluorescence microscopy for histone 3 lysine 36 dimethylation (H3K36me2) was performed. n=3, t-test. **C)** 0082T CAFs were cultured in 1% O_2_ for 120 hr, and iterative indirect immunofluorescence imaging (4i) was performed for the indicated proteins. n=3, one-way ANOVA and Games-Howell post-hoc test. **D)** and **E)** 0082T CAFs were engineered for stable expression of two different shRNAs for NSD2 knockdown or a control shRNA. The cells were grown for 120 hr at 1% O_2_, and immunofluorescence microscopy was performed for HGF (D) or IGF-2 (E). n=4, one-way ANOVA and Tukey multiple comparison test. **F)** 4i was performed on KPCY tumor sections to quantify NF-κB and H3K36me2 expression in CAFs. CAFs were identified as cells positive for podoplanin (PDPN) and negative for the YFP epithelial lineage tracer. CAFs were also assessed for their staining for Hypoxyprobe (HYP). n=4, t-test. Data are presented as mean ± SEM for bar plots (A, D, E), as violin plots overlayed with box-and-whisker plots that represent the first quartile (lower box bound), median (center bound), or third quartile (upper box bound) (B, C), or as a box-and-whisker plot with individual data points shown (F). * *p* <0.05, ** *p* < 0.01, *** *p* < 0.001, **** *p* < 0.0001

Given the lack of effect of HIF knockdown, we shifted focus to histone demethylases that exhibit reduced activities in hypoxic conditions (*19, 30, 31*). The reduced activity of lysine demethylase KDM2A in hypoxia promotes a PDAC cancer cell EMT that is dependent on histone 3 lysine 36 dimethylation (H3K36me2) (*19*). Additionally, hypoxia reduces the activities of KDM5A and KDM6A, resulting in increased histone 3 lysine 4 trimethylation (H3K4me3) and histone 3 lysine 27 trimethylation (H3K27me3) (*30, 31*). In 0082T CAFs, hypoxia increased H3K36me2 (**Figure 5B**) and H3K4me3 (**Supp Figure S9A**) but decreased H3K27me3 (**Supp Figure S9B**). A KDM5 inhibitor did not affect the expression of HGF in hypoxic CAFs and was therefore not tested further (**Supp Figure S9C**).

H3K36me2 increases the expression of some NF-κB target genes (*67*), and KDM2A can demethylate p65 to impair NF-κB binding to DNA (*68*). Therefore, NF-κB transcriptional activity may be enhanced by hypoxia. Because of the reported roles of H3K36me2 and KDM2A on NF-κB pathway activity, we focused on H3K36me2 in further experiments and postulated that H3K36me2 cooperates with NF-κB to promote IGF-2 and HGF expression in hypoxic CAFs. To test this, the abundance of H3K36me2, NF-κB, IGF-2, and HGF were assessed in hypoxic 0082T CAFs using iterative indirect immunofluorescence imaging (4i). 4i enabled this experiment on a typical four-color microscope with antibodies that would otherwise have been incompatible (see *Methods*). Analysis of images demonstrated that CAFs with high nuclear NF-κB abundance and high H3K36me2 displayed greater expression of IGF-2 and HGF than CAFs with high nuclear NF-κB or high H3K36me2 alone (**Figure 5C**). To test the importance of H3K36me2 more directly, we transduced 0082T CAFs with vectors encoding shRNAs for NSD2 knockdown, which decreased NSD2 and H3K36me2 signals (**Supp Figure S9D**). As hypothesized, NSD2 knockdown diminished expression of HGF and IGF-2 (**Figure 5D, E**). Collectively, these results indicate that nuclear NF-κB and H3K36me2 were increased in CAFs in response to hypoxia and cooperated to promote the expression of HGF and IGF-2. To extend this analysis *in vivo,* we performed 4i on KPCY tumor sections. Hypoxic CAFs displayed significantly higher nuclear accumulation of NF-κB and nuclear H3K36me2 than their normoxic counterparts (**Figure 5F**). Interestingly, 20107 cells (iCAF-like) exhibited higher nuclear translocation of NF-κB but lower H3K36me2 than 15760 cells (myCAF-like) (**Supp Figure S9E, F**). Thus, the relevance of the hypoxia-mediated mechanisms we uncovered for stable iCAF-like phenotypes is unclear.

Hypoxic tumor regions contain numerous interfaces between malignant cells and CAFs, and juxtacrine interactions with cancer cells can activate CAF signaling pathways (*69*). To test for juxtacrine regulation of NF-κB and H3K36me2 in hypoxic CAFs, we used a co-culture approach with fixed cancer cells. GFP-labelled HPAF-II cells were maintained at 1% O_2_ for 120 hr and fixed to preserve membrane proteins and prevent the release of secreted factors. 0082T CAFs were then cultured with fixed HPAF-II cells for 48 hr. H3K36me2 and nuclear NF-κB were elevated in the 0082T cells near (< 20 μm) HPAF-II cells in both 21% and 1% O_2_ conditions (**Supp Figure S10A, B**). NF-κB was elevated in hypoxic 0082T cells (near and far from HPAF-II) compared to normoxic CAFs. H3K36me2 was not significantly different between 21% and 1% O_2_, which may be due to the 0082T being reoxygenated during seeding onto plates with fixed HPAF-II cells and limited hypoxia exposure time (48 hr). That issue notwithstanding, these results indicate that CAF-malignant cell juxtacrine interactions may have cooperated with cell-autonomous hypoxia effects to enhance histone methylation and NF-κB activation in CAFs.

### Hypoxic CAFs reside proximal to hypoxic mesenchymal cancer cells in PDAC tumors

To explore the ability of hypoxic CAFs to promote EMT in tumors, we performed 4i on sections of KPCY tumors with staining for podoplanin (PDPN), Hypoxyprobe (HYP), YFP (lineage-traced cancer cells), and vimentin (**Figure 6A**). The multiplexed micrographs were used to classify cells as CAFs (PDPN+) or cancer cells (YFP+). CAFs were further sub-classified as normoxic (HYP-) or hypoxic (HYP+). Similarly, cancer cells were sub-classified as normoxic or hypoxic and vimentin-positive (Vim+) or -negative (Vim-) (**Figure 6B**). Importantly, hypoxic CAFs were typically located near hypoxic vimentin-positive cancer cells (**Figure 6B**). Using the classifications and spatial coordinates for each cell, distances were computed between all CAFs and hypoxic cancer cells. Hypoxic cancer cells were binned according to the most common CAF subtype (HYP-and HYP+ CAFs) within a 50-μm radius (see *Methods*). The aggregated analysis indicated that hypoxic cancer cells near hypoxic CAFs displayed more vimentin expression than those farther from hypoxic CAFs (**Figure 6C**). Thus, hypoxic CAFs and hypoxic mesenchymal cancer cells tended to be co-located within a length scale over which paracrine signaling interactions are estimated to occur (∼100 μm) (*70, 71*).

**Figure 6.**
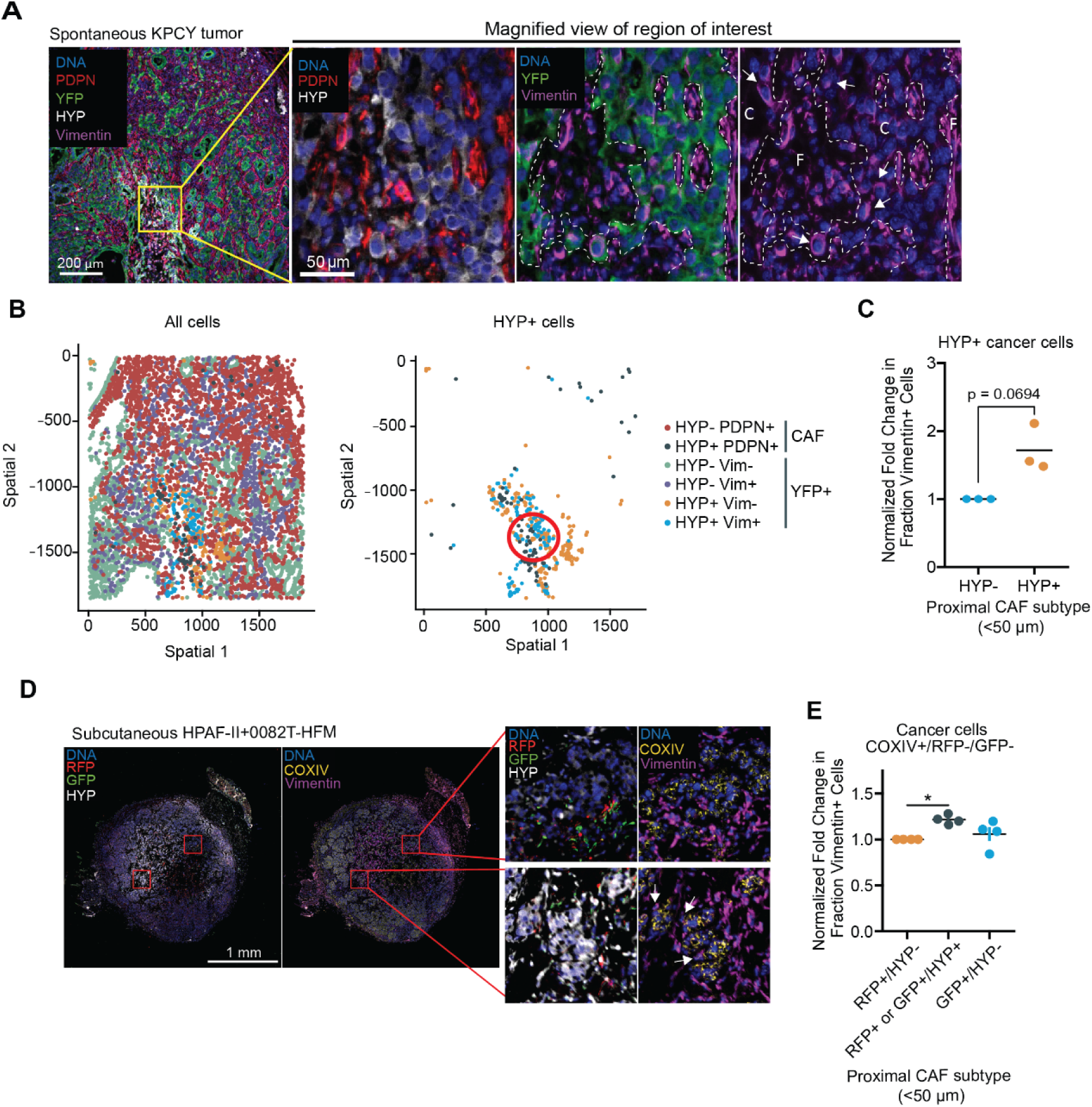
Mesenchymal cancer cells are found near hypoxic CAFs in PDAC mouse models. **A)** Indirect iterative immunofluorescence imaging (4i) was performed on KPCY tumor sections for the indicated proteins. CAFs were identified as cells positive for podoplanin (PDPN+) and negative for the epithelial lineage tracer YFP (YFP-). Staining for vimentin and Hypoxyprobe (HYP) was also performed. White dashed outlines in the image insets separate cancer cells (“C”) from fibroblasts (“F”). White arrows indicate vimentin-positive cancer cells. Image representative of n=3. **B)** In KPCY tumor section images, CAFs and cancer cells were classified based on marker expression and identified as normoxic CAFs (HYP-/PDPN+), hypoxic CAFs (HYP+/PDPN+), normoxic vimentin-negative cancer cells (HYP-/Vim-/YFP+), normoxic vimentin-positive cancer cells (HYP-/Vim+/YFP+), hypoxic vimentin-negative cancer cells (HYP+/Vim-/YFP+), or hypoxic vimentin-positive cancer cells (HYP+/Vim+/YFP+). The red circle in HYP+ cells spatial map indicates an area that was enriched with hypoxic CAFs and hypoxic vimentin-positive cancer cells. **C)** Hypoxic cancer cells (HYP+) were binned based on proximity to hypoxic CAFs. The fraction of vimentin-positive hypoxic cancer cells was compared between cells that were near hypoxic CAFs (<50 μm) or far from them (>50 μm). Data were normalized for each tumor to the “far” result to obtain fold-change for the fraction vimentin-positive cells. n=4, paired t-test. **D)** 4i for the indicated proteins was performed on sections of subcutaneous tumors generated in mice using HPAF-II cells and 0082T CAFs engineered with a hypoxia fate mapper reporter (0082T-HFM). **E)** For the tumors described in panel (D), CAFs were classified as hypoxic (HYP+), normoxic (HYP-), or once-hypoxic (GFP+/HYP-). Cancer cells were identified using human-specific COXIV staining and binned as vimentin-negative or - positive. Cancer cells were identified by expression of the human cell marker COXIV and categorized as being near normoxic CAFs (HYP-), hypoxic CAFs (HYP+), or once hypoxic CAFs (GFP+/HYP-). The fraction vimentin-positive cancer cells were normalized to those near normoxic CAFs for each tumor to obtain fold-change in fraction vimentin-positive for each sample. n=4, one-way ANOVA with Tukey’s multiple comparison test. Data are presented as mean (horizontal line) with data points shown for each biological replicate (C, E). * *p* <0.05, ** *p* < 0.01, *** *p* < 0.001, **** *p* < 0.0001

### Once-hypoxic CAFs do not co-locate with mesenchymal cancer cells in tumors

Hypoxia can promote durable cancer cell EMT, ROS-resistance, and glycolytic phenotypes that persist after reoxygenation (*19, 29, 72*). To test if the effect of hypoxia on CAFs was durable, we engineered 0082T CAFs with a hypoxia fate-mapping reporter that causes cells to stably convert from DsRed to GFP expression following sufficient exposure to low oxygen (*72*) and selected a clonal population with the desired switching characteristics (**Supp Figure S11A**). Subcutaneous tumor xenograft experiments were performed in mice using HPAF-II cells and engineered hypoxia fate-mapping 0082T cells (0082T-HFM). Because human CAFs are rapidly depleted in tumor xenografts (*73*), we limited tumor outgrowth to 7 and 14 days and found that 0082T-HFM CAFs were detected in tumors at the day-7 time point only (**Supp Figure S11B**). Sections of day-7 tumors were subjected to 4i, and individual cells were classified based on marker expression (**Figure 6D**). Cancer cells were identified based on expression of human COXIV and lack of RFP or GFP (COXIV+/RFP-/GFP-), and subclassified as vimentin-positive or -negative. CAFs were classified as never-hypoxic (RFP+/HYP-), hypoxic (RFP+/HYP+ or GFP+/HYP+), or once-hypoxic (GFP+/HYP-). Cancer cells were then binned as being near one of the three CAF subtypes based on the one most frequently encountered within 50-μm (*Methods*). Compared to cancer cells near never-hypoxic CAFs, cancer cells near hypoxic CAFs displayed more frequent vimentin expression, but those near once-hypoxic CAFs displayed no difference (**Figure 6E**). This suggests that the enhanced ability of hypoxic CAFs to drive EMT was not a durable phenotype following hypoxia exposure and subsequent reoxygenation. Indeed, CM from hypoxic 0082T CAFs that were reoxygenated for 24 hr prior to conditioning medium produced fewer vimentin-positive HPAF-II cells than CM from 0082T cells that were maintained continuously in hypoxia (**Supp Figure S11C**).

## DISCUSSION

We identified a hypoxia-regulated mechanism that converts CAFs to iCAF-like cells that secrete EMT-promoting ligands. The mechanism is strong enough to cooperate with hypoxia-mediated autonomous EMT in cancer cells. In our proposed mechanism (**Figure 7**), which bares the characteristic multiscale hallmarks of other tumor microenvironment-regulated and cancer cell signaling phenomena (*57, 74*), hypoxia drives NF-κB activation in CAFs through ROS accumulation and a concomitant increase in H3K36me2, which likely occurs due to impaired KDM2A activity in hypoxia (*19*). The cooperation of these effects in hypoxic CAFs promotes expression of HGF and IGF-2, which drive cancer cell EMT. Interestingly, H3K36me2/3 supports NF-κB target gene transcription in other settings (*75*), suggesting that the pathway we uncovered may be a generally applicable hypoxic mechanism for transcriptomic rewiring. In tumors, hypoxic CAFs and hypoxic mesenchymal cancer cells were preferentially located within a length scale characteristic of paracrine signaling interactions, but this spatial relationship was lost for once-hypoxic CAFs in normoxic tissue.

**Figure 7.**
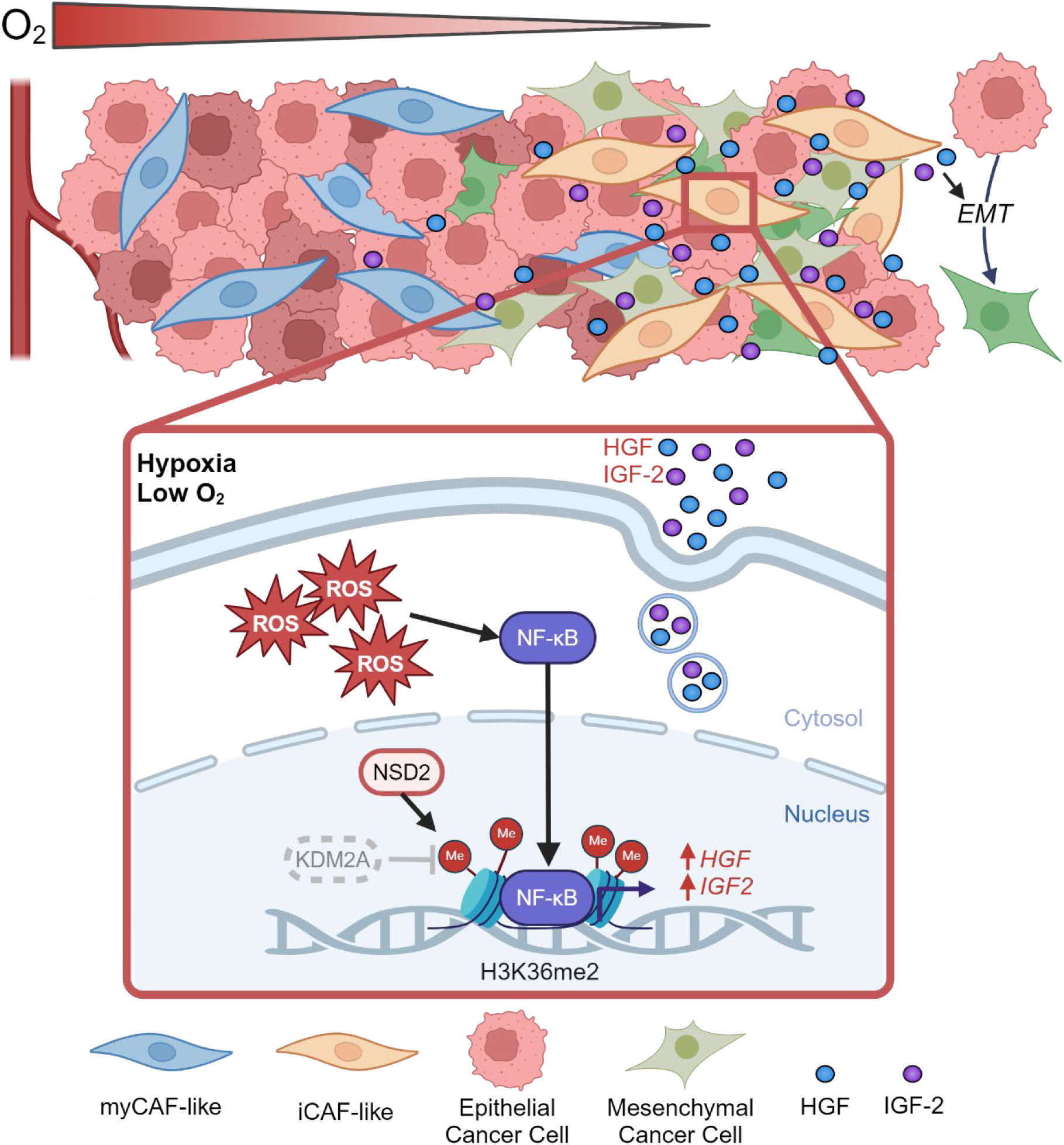
Hypoxia promotes expression of EMT-inducing growth factors by CAFs through regulation of histone methylation and NF-κB signaling. In the mechanism we propose in hypoxic CAFs, reactive oxygen species (ROS) promote NF-κB nuclear translocation. A concomitant reduction in the activity of histone demethylase KDM2A promotes dimethylation of histone 3 at lysine 36 (H3K36me2) and cooperates with NF-κB nuclear localization, potentially by exposing promoter binding sites, to enhance HGF and IGF-2 expression. CAF-secreted HGF and IGF-2 cooperate to drive EMT in PDAC cancer cells.

Our finding on the role of H3K36me2 in CAFs extends understanding of the importance of this epigenetic mark to new PDAC cell types. Previously, the role for H3K36me2 in PDAC had been studied only in cancer cells, where it is required for a hypoxia-mediated cell-autonomous EMT (*19*) and TGFβ-driven EMT (*76*). Our findings motivate the continued development of inhibitors targeting histone 3 lysine 36 methyltransferases, which could simultaneously target EMT-driving processes in ductal cells and CAFs. While we focused on H3K36me2, other histone methylation sites, including histone 3 lysine 4 and histone 3 lysine 27, also undergo methylation in response to hypoxia due to diminished histone demethylase activities (*30, 31, 76*). Other hypoxia-regulated histone modifications, including acetylation (*77*), may also inform PDAC CAF subtype switching. Thus, histone deacetylases may be useful therapeutic targets in PDAC as they promote pro-inflammatory pathways in CAFs and expression of iCAF marker genes (*5*).

While our study focused on the impact of hypoxia on CAF cytokine secretion, hypoxia may also induce CAF metabolic changes that impact EMT or otherwise affect disease progression. Indeed, we observed that hypoxia augmented CAF LDHA expression. Although we concluded that LDHA-generated lactate was unlikely to be responsible for ROS-dependent NF-κB activity or growth factor expression, LDHA-generated lactate from hypoxic CAFs could have other important functions. In PDAC, lactate inhibits immune responses by recruiting myeloid-derived suppressor cells and consequently reducing the efficacy of radiotherapy (*8*). In breast cancer, lactate produced by hypoxic CAFs supports cancer cell biosynthesis (*29*).

Our investigations focused on CAF-cancer cell paracrine interactions, but CAF-secreted cytokines may regulate other important cell types in tumors. Hypoxic CAFs promote an immunosuppressive M2 macrophage polarization in a HIF2α-dependent manner (*28*), which may arise through iCAF-secreted CXCL12 (*49*). Interestingly, M2 macrophages have also been linked to EMT and metastasis in PDAC through CCL18 (*78*) and IL-8 (*79*), suggesting that hypoxic CAFs could also facilitate a macrophage-mediated and EMT-driving cell crosstalk. Furthermore, hypoxic CAFs generate angiogenic cytokines [e.g., VEGF in breast cancer (85)], which may modulate tumor oxygenation and related cell phenotypes.

Determining the impact of chemotherapy and radiation on CAF heterogeneity will also be important for understanding how to target CAF-mediated cell crosstalk effectively. PDAC patients treated with gemcitabine or FOLFIRINOX display a higher proportion of iCAFs than myCAFs (*80*), but it is unknown if this skew results from phenotype switching or preferential subtype susceptibility to chemotherapy. Given that hypoxia reduces cell proliferation and that chemotherapy primarily affects rapidly dividing cells, it is possible that hypoxic CAFs are simply more resistant to chemotherapy, resulting in iCAF enrichment following treatment.

Our results also indicated a lack of HIF1α and HIF2α involvement in HGF and IGF-2 secretion by hypoxic CAFs. These results were surprising given that HIF1α was reported to be required for conversion to an iCAF phenotype in murine PSCs (*44*). Thus, it is possible that the hypoxia-induced iCAF phenotype has HIF-dependent and -independent characteristics. There may also be differences between how murine- and human-derived PSCs/CAFs behave in hypoxic environments that are yet to be uncovered.

While our studies primarily focused on the effects of hypoxia on isolated CAFs, we also found that cancer cell EMT was more prominently induced by CM from CAF-cancer cell co-cultures than from CAF monocultures. The augmented effect of co-culture CM may have resulted from hypoxic cancer cells reinforcing the iCAF phenotype through their reported ability to secrete IL1α in hypoxic conditions (*43*). Our finding that juxtracrine signaling between CAFs and cancer cells promoted nuclear NF-κB accumulation and H3K36me2 in CAFs, which we showed promotes the secretion of HGF and IGF-2, may also be at play. Collectively, these results highlight the importance of CAF-cancer cell crosstalk in generating the iCAF-like cells that promote cancer cell EMT.

The effects of hypoxia on CAF phenotype we investigated were reversible, but it is worth recalling that the 20107 (iCAF) and 15760 (myCAF) CAF cell lines isolated from human PDAC tumors were essentially stuck in iCAF- or myCAF-like states, respectively. It is possible that the diverse origins of CAFs may predispose some subgroups to phenotypic plasticity and others to specific CAF subtypes, such as those observed for mesothelium-derived antigen-presenting CAFs (*81*). 20107 and 15760 CAF cells have not been sequenced, but it is also possible that acquired mutations or other therapy-induced alterations contributed to the durable retention of iCAF characteristics in 20107 cells. Hypoxia can regulate epigenetic modifications and promote durable alterations to cellular phenotypes (*29*). Indeed, hypoxia induces a durable EMT in PDAC cancer cells (*19*), and hypoxic breast cancer cells acquire a ROS-resistant phenotype that persists following reoxygenation (*72*). It is possible that the hysteretic effect of hypoxia in those cancer cell settings requires the cooperation of an oncogene not present in CAFs.

## METHODS

### Cell culture

HPAF-II human PDAC cells (Carl June, University of Pennsylvania) were maintained in RPMI medium (ThermoFisher: #11875093) supplemented with 10% fetal bovine serum (FBS; VWR: 97068-085), 1 mM L-glutamine (ThermoFisher: #25030081), 100 units/mL penicillin (ThermoFisher: #15140122), and 100 μg/mL streptomycin (ThermoFisher: #15140122). Capan-2 human PDAC cells (ATCC) were maintained in DMEM (ThermoFisher: #11-965-092) with 10% FBS, 1 mM L-glutamine, 100 units/mL penicillin, and 100 μg/mL streptomycin. 0082T CAF cells derived from human PDAC tumors and immortalized with SV40 large T antigen were created by Dr. Andrew Lowy and shared by Dr. Mara Sherman (*82*). 0082T cells were maintained in DMEM with 10% FBS, 1 mM L-glutamine, 100 units/mL penicillin, and 100 μg/mL streptomycin. Primary PSCs (ScienCell) were cultured in pancreatic stellate cell medium (ScienCell: #5301) following manufacturer recommendations. Primary human CAF lines ST-00020107 (20107) and ST-00015760 (15760) were maintained in F medium, a 3:1 mix of DMEM and F12 (ThermoFisher: #11765054), supplemented with 5% FBS, 10 ng/mL EGF (Peprotech: #AF-100-15), 24 μg/mL adenine (Sigma: #A8626), 5 μg/mL insulin (Sigma: #I5500), 0.4 μg/mL hydrocortisone (Sigma: #H0888), 10 μM Y-27632 (Selleck Chem: #S6390), and 8.4 ng/mL cholera toxin (Sigma: #C8052). Media for 20107 and 15760 cells was replaced with complete DMEM for conditioning experiments to maintain consistency with other conditioned medium experiments and avoid the potentially confounding effects of the supplements in complete F medium. Cell lines were tested for mycoplasma using the MycoAlert PLUS detection kit (Lonza: #LT07-710). HPAF-II cells were authenticated by short tandem repeat profiling using GenePrint 10 (Promega) by the Genetic Resources Core Facility at the John Hopkins University School of Medicine.

### Conditioned medium

0082T CAFs, PSCs, and primary human CAFs (20107 and 15760) were plated at 1×10^6^ cells per 10-cm cell culture plate in their standard media. One day after plating, cells were either maintained under normal culture conditions (i.e., 21% O_2_) with fresh DMEM or switched to various experimental conditions (i.e., 1% O_2_ with or without added inhibitors). 72 hr later, cell culture medium was replaced with fresh complete DMEM (without inhibitors) for all cell types, and cells were returned to their respective oxygen conditions for 48 hr to condition medium prior to its collection. The removal of any inhibitors for the 48 hr conditioning period prevented carryover to cells treated with conditioned medium (CM). For experiments involving reoxygenated 0082T CM, 0082T CAFs were cultured in 1% O_2_ for 120 hr with media changes at 24 hr and 72 hr before replating at 2×10^6^ cells in a 10 cm plate and moving the cells to 21% O_2_.

After 24 hr, the media was replaced on the reoxygenated 0082T cells and conditioned for 48 hr before collecting as CM. All CM was passed through a 0.22-μm syringe filter to remove cell debris. CM samples were diluted 1:1 with complete RPMI or DMEM (for treatment of HPAF-II or Capan-2 cells, respectively) and aliquoted prior to freezing at −80°C for future use. HPAF-II or Capan-2 cells were treated with CM at 24 and 72 hr after initial seeding onto plates for experiments. For experiments utilizing molecular weight filters, undilute 0082T CM was loaded into 3- and 30-kDa ultrafilters and centrifuged for 30 min at 4,000 rcf. Fractions were diluted with complete RPMI before using CM fractions to treat HPAF-II cells as previously described.

### Growth factors and inhibitors

Recombinant IL1α, TGFβ1, IGF-2, and HGF (all from Peprotech) were used at 10 ng/mL, 10 ng/mL, 100 ng/mL, and 10 ng/mL, respectively. All inhibitors were reconstituted in DMSO, and concentrations were chosen based on prior studies as referenced. IKK-16 (Selleck) was used at 0.5 μM (*83*). PHA665752 (Santa Cruz Biotechnology) and linsitinib (Selleck) were used at 2.5 μM (*84, 85*). Stattic (Abcam) and CI-1040 (LC Laboratories) were used at 1 μM (*19, 86*). GSK2837808A (MedChemExpress) and galunisertib (Selleck) were used at 10 (*19, 87*). CAFs or PSCs were pretreated (before conditioning media) with specific growth factors and inhibitors diluted in complete DMEM (for all cell types) for 72 hr. For cancer cell EMT studies, growth factors or inhibitors were diluted in RPMI and replenished every 48 hr.

### Kinase inhibitor screen

HPAF-II cells were plated in 96-well format. One day after plating, six control wells were treated with DMSO (vehicle) to measure baseline vimentin expression. Another six wells were treated with CM from hypoxic 0082T cells plus DMSO to determine the baseline ability of CM to augment vimentin expression. A library of 430 small-molecule inhibitors (Selleck: Library-Z188031-100uL-L1200) was diluted from 1 mM stocks to 20 μM using DMSO. The inhibitor library was stored in 96-well v-bottom polypropylene plates (Corning: #3343) with aluminum plate coverings (ThermoFisher: #AB0718) at −80°C. For non-control wells, HPAF-II cells were treated with CM and 10 μL of each inhibitor, with mixing by pipetting, to achieve a 1 μM working concentration for each inhibitor. 1 μM was chosen to minimize cell toxicity while still achieving a phenotypic effect. Treatments with DMSO, CM alone, or CM plus inhibitor were repeated after 72 hr. 48 hr later (120 hr after first treatment), cells were fixed and stained for immunofluorescence imaging using antibodies against vimentin and ERK (the latter for cell segmentation). Percent-inhibition of vimentin expression was calculated for each compound using DMSO-only and CM+DMSO as the minimum and maximum vimentin expression values, respectively. Inhibitors were classified as effective if they reduced the fraction of vimentin-positive cells by >50% compared to the CM+DMSO condition.

### Immunofluorescence microscopy

To quantify EMT and signaling pathway activation among cells, immunofluorescence microscopy was frequently used in this study because of its utility in assessing heterogeneity within cell populations (*88*). Cells were fixed using 4% paraformaldehyde (PFA) (Thomas Scientific: #C994J69) in PBS for 20 min. Following fixation, cells were permeabilized with 0.25% Triton-X100 (Sigma: #X100-100ML) in PBS (ThermoFisher: #14-190-144) for 5 min. Cells were then stained using primary antibodies diluted in intercept blocking buffer (IBB; VWR: #927-60001) overnight at 4°C. Antibody dilutions for immunofluorescence microscopy are provided in **Supplementary Table S1**. Samples were then washed with Tween-20 (Fisher: #BP337-100) in PBS five times before incubating with secondary antibodies (1:750) and Hoechst (1:2000) diluted in IBB for 1 hr at 37°C. For experiments on coverslips, samples were mounted on glass slides using Prolong Gold Antifade (Fisher: #P36930) before imaging. For 96-well plates, a solution of 20% glycerol in PBS was added to each well for imaging. Cells were imaged using 10× or 20× objectives on a Zeiss Axiovert Observer.Z1 fluorescence microscope or BioTek Cytation5 imaging plate reader. All comparisons were made across images with identical exposure times and image settings. At least four frames were taken randomly for each biological replicate to image a total of at least 1000 cells.

### Iterative indirect immunofluorescence imaging (4i) of 0082T cells on tissue culture plastic

The 4i protocol and buffers were used and prepared as described in Gut et al. (*89*), with the exception of the blocking buffer which we formulated using IBB +150 mM maleimide. 0082T CAFs were fixed with 4% PFA for 20 min and then permeabilized using 0.25% Triton-X100 in PBS for 5 min. Once permeabilized, samples were incubated with 4i blocking buffer. Primary antibodies were diluted in IBB at manufacturer-recommended concentrations and incubated overnight at 4°C. Samples were washed with 1× PBS 5 times. Samples were then incubated with Hoechst (1:2000) and appropriate secondary antibodies diluted in IBB (1:750) for 1 hr at room temperature. Samples were washed with DI water 5 times before adding 4i imaging buffer and then imaged. For each round, imaging was performed as described in *Immunofluorescence microscopy*. Antibodies were eluted by washing the plate with 4i elution buffer 3 times followed by washing with PBS 5 times. Hoechst diluted in PBS was added to samples for 5 min for post-elution imaging to ensure antibody removal occurred. Samples were then washed with DI water 5 times. Following post-elution imaging, samples were blocked and re-stained with different antibodies using the workflow already described. The complete list of antibodies by round of 4i staining for 0082T cells (**Figure 5C**) is provided in **Supplementary Table S2.** Images from the two rounds of staining were aligned using the CellProfiler *Align* module with the nuclear stain serving as the reference for image alignment.

### Automated image analysis for fluorescence microscopy

For fixed-cell immunofluorescence images, background autofluorescence was reduced using the Gen5 (BioTek) rolling-ball background subtraction data reduction protocol. CellProfiler was used to identify individual cells using DNA (Hoechst stain) as a primary object (nuclear segmentation marker). Cell segmentation for fixed-cell immunofluorescence was performed using E-cadherin or ERK as a secondary cytoplasmic signal. Cells were classified as positive for a marker (e.g., vimentin) using a threshold based on a secondary antibody-only control that was applied to all images. While most fixed-cell signals were quantified on a whole-cell basis, E-cadherin was quantified at cell edges, H3K36me2 was quantified in nuclei, and NF-κB was quantified as a nucleus-to-cytoplasm ratio.

For tumor section images, the background signal was removed by subtracting the bottom 10% of image intensity by pixel using the *Image Math* module in CellProfiler. The nucleus was used as the primary object, and podoplanin, COXIV, or YFP signal was used as a secondary cytoplasmic signal. Thresholds used for classifying cell types (positive or negative for a given marker) were specified separately for each tumor and adjusted manually by inspection of morphology markers in images to ensure accurate identification of cell types and states. Because CellProfiler struggled to quantify cytoplasmic NF-κB in tumors, nuclear NF-κB signal (rather than the nucleus-to-cytoplasm ratio) was reported for those samples.

### Genetically engineered mouse models

Kras^LSL-G12D^, p53^LSL-R172H^, Pdx1-Cre, Rosa26^LSL-YFP^ (KPCY) mice were described previously (*90*). Both male and female mice were used. Mice were examined for signs of morbidity twice per week. Mice were injected with 60 mg/kg pimonidazole (referred to as “Hypoxyprobe” throughout; Hypoxyprobe: #HP7-100) 90 min prior to sacrifice. Tissue samples were fixed overnight in zinc-buffered formalin and then paraffin-embedded the following day. Animal procedures and experiments were conducted in accordance with NIH guidelines for animal research and approved by the University of Pennsylvania Institutional Animal Care and Use Committee.

### Hypoxia fate-mapping cell line engineering

0082T cells were engineered with a previously described reporter that causes cells to switch from DsRed to GFP expression irreversibly in response to hypoxia (*72*). LX293T cells were transfected with the CMV-loxp-DsRed-loxp-eGFP (Addgene: #141148) or 4xHRE-MinTK-CRE-ODD (Addgene: #141147) plasmid and the psPAX2 (Addgene: #12260) and pMD2.G (Addgene: #12259) packaging plasmids. CMV-loxp-DsRed-loxp-eGFP and MinTK-CRE-ODD were provided by Dr. Daniele Gilkes (Johns Hopkins). psPAX2 and pMD2.G plasmids were provided by Dr. Didier Trono. Polyjet (SignaGen: #SL100688) was used to improve transfection efficiency. 0082T cells were initially transduced with filtered (0.45 µm) viral supernatant from LX293T cells transfected with CMV-loxp-DsRed-loxp-eGFP and 8 μg/mL of polybrene. 0082T cells were then selected using zeocin (400 μg/mL) for 10 days. Following selection, 0082T CAFs were transduced with filtered viral supernatant from LX293T cells transfected with 4xHRE-MinTK-CRE-ODD with 8 μg/mL of polybrene. Transduced 0082T CAFs were cultured in 5% O_2_ and single-cell sorted into 96-well plates to retain DsRed+ cells and remove GFP+ cells using a Sony MA900 Cell Sorter by the UVA Flow Cytometry Core Facility, with parental 0082T CAFs used to set gates. The viability of sorted clones was improved by using complete medium conditioned by parental 0082T CAFs supplemented with an additional 10% FBS. Once wells were confluent, each 96-well plate produced in the sorting step was split into three sub-cultured plates to create three copies of each surviving clone, with one plate maintained at 21% O_2_, one at 3% O_2_, and one at 1% O_2_. Clones were screened using a Cytation5 imaging plate reader (BioTek) with a 10× objective to identify those that exhibited increased GFP expression in 1% O_2_ only. Only one clone met those criteria, which we refer to as hypoxia fate-mapping 0082T cells (0082T-HFM).

### Subcutaneous tumors containing 0082T hypoxia fate-mapped cells

NOD/SCID/IL2Rγ (NSG) mice, purchased from Jackson Laboratory (Strain: #005557, Jackson Laboratory, Bar Harbor, ME), were injected subcutaneously with 1×10^6^ HPAF-II and 2×10^6^ 0082T-HFM cells suspended in 60 μL of Matrigel (Fisher Scientific: #CB-40234C). Tumors were allowed to grow for 7 or 14 days. Mice were injected with 60 mg/kg Hypoxyprobe reagent 90 min prior to sacrifice. Tissue samples were fixed using zinc formalin and embedded in paraffin. Animal procedures and experiments were conducted in accordance with NIH guidelines for animal research and approved by the University of Virginia Institutional Animal Care and Use Committee.

### Antigen retrieval

Formalin-fixed paraffin-embedded (FFPE) slides were cut to a thickness of 5 µm by the UVA Research Histology Core. FFPE slides were deparaffinized by two consecutive 10-min xylene washes. Slides were then submerged in 100% EtOH, then 90% EtOH, and finally 80% EtOH, with a 5-min incubation for each. Following EtOH washes, slides were washed with deionized (DI) water before antigen retrieval (AR). AR buffer was made using 1.2 g Tris base, 2 mL 0.5 M EDTA-pH 8.0, Tween-20 (0.05%), and DI water to 1L. The pH of the AR buffer was adjusted to 9 using HCl (Sigma: #HX0603S). AR buffer was heated to 95°C before submerging slides for 20 min. Following AR, slides were washed with DI water three times and used for immunofluorescence staining or iterative indirect immunofluorescence imaging.

### 4i for tumor sections

FFPE samples were antigen-retrieved at pH 9, as previously described. Samples were then permeabilized with 0.1% Triton-X in PBS for 20 min at room temperature. Blocking, primary antibody incubation, and wash steps were conducted as described earlier for 0082T cells. Following primary antibody incubation and subsequent wash steps, samples were incubated with Hoechst (1:1000) and appropriate secondary antibodies diluted in IBB (1:500) for 1 hr at room temperature. For the imaging step, 10% glycerol was added to the 4i imaging buffer to facilitate removal of glass coverslips following imaging. Glass coverslips were carefully placed on each slide and inspected to ensure the absence of bubbles prior to imaging. Following imaging, glass coverslips were carefully removed by floating slides in water until they detached. Samples were eluted with 4i elution buffer and washed as previously described. Samples were then stained with Hoechst for post-elution imaging, as described previously, and imaged. This protocol was repeated until all antibody signals were imaged. The 4i approach for assessing NF-κB and H3K36me2 in KPCY sections (**Figure 5F**) is provided in **Supplementary Table S3**. The list of antibodies by round for 4i staining of KPCY tumors (**Figure 6A**) is provided in **Supplementary Table S4**. The 4i staining approach for subcutaneous tumors containing HPAF-II and 0082T-HFM cells (**Figure 6D**) is provided in **Supplementary Table S5**.

### Spatial analysis of tumor-section 4i

Automated image analysis was performed using CellProfiler (*91*). Images from three rounds of staining were aligned using the CellProfiler *Align* module with the nuclear stain serving as the reference for each round. Aligned images were then used for downstream analysis. Individual cells were identified using the Hoechst nuclear stain. Nuclei were identified as primary objects and were used to measure all nuclear signals. All other stains were identified as secondary objects. The *Measure Object Intensity* module was used to quantify signals for each analyte. Secondary antibody-only controls were used to set signal cutoffs to determine whether cells were positive for specific signals. Cell classifications were determined based on staining for specific markers. Podoplanin and GFP were used to identify CAFs and cancer cells, respectively. Hypoxyprobe staining was used to subclassify CAFs and cancer cells as hypoxic (HYP+) or normoxic (HYP-). Vimentin was used as a mesenchymal marker to classify cancer cells further. Cell coordinates were exported along with single-cell fluorescence intensities.

CellProfiler coordinates, exported as pixel units, were converted to microns based on image resolution. Using the *Fields* R package, distances between all pairs of cell types of interest were computed using the *rdist* function based on the centroids of each nucleus. For KPCY tumors, hypoxic cancer cells (HYP+/YFP+) were binned based on the most common hypoxia status (HYP- or HYP+) of CAFs within a 50-μm radius. The fraction of vimentin-positive cancer cells was then determined for both cancer cell bins, and normalization was performed within each tumor to the result for cancer cells proximal to HYP-CAFs to calculate fold-changes in vimentin-positivity. For subcutaneous tumors formed using HPAF-II and 0082T-HFM cells, HPAF-II cells were binned based on the most common status (RFP+/HYP-, RFP+ or GFP+/HYP+, and GFP+/HYP-) of CAFs within a 50-μm radius. The fraction of vimentin-positive cancer cells was then determined for all three cancer cell bins, and normalization was performed within each tumor to the result for cancer cells proximal to RFP+/HYP- CAFs.

### Quantitative real-time PCR

RNA was isolated from cells using the RNeasy Kit (Qiagen: #74104) and reverse-transcribed using the High-Capacity cDNA Reverse Transcription Kit (Applied Biosciences: #4368814). PowerUp SYBR Green (Applied Biosciences: #4367659) detection was used for qRT-PCR. Relative changes in transcript abundances were determined using the ΔΔCt method. Gene fold-change data were normalized using *CASC3* for hypoxia experiments or *GAPDH* for experiments in oxygenated conditions. Primer sequences are provided in **Supplementary Table S6**.

### Western blotting

Cell lysates were prepared using a cell extraction buffer (Invitrogen: #FNN0011) supplemented with phosphatase and protease inhibitors (Sigma-Aldrich: #P8340, #P0044, and #P5726). Crude lysates were centrifuged for 10 min at 18,000 rcf to obtain clarified fractions. A micro-bicinchoninic acid (BCA) assay (Pierce; Thermofisher: #23225) was used to measure protein concentrations. 20 μg of protein were mixed with 10× NuPAGE reducing agent (Invitrogen: #NP0005), 4× LDS sample buffer (Invitrogen: NP0007), and ultrapure water. Samples were boiled at 100°C for 10 min and run on a 1.5 mm NuPAGE gradient (4-12% gradient) gel (Invitrogen: #NP0336BOX). Gels were transferred to a 0.2 μm nitrocellulose membrane (BioRad: #1704271) using a BioRad TurboBlot system. Membranes were blocked using IBB for 1 hr on an orbital shaker. Primary antibodies were diluted in IBB at manufacturer-recommended concentrations and incubated on the membrane overnight at 4°C with shaking. The following day, membranes underwent three 5-min washes using 0.1% Tween-20 in PBS on an orbital shaker. Secondary antibodies were diluted in IBB (1:10,000) and incubated for 2 hr with shaking at room temperature. Membranes were washed as described previously with 0.1% Tween-20 in PBS. Membranes were imaged using a LI-COR Odyssey CLx.

### siRNA-mediated knockdowns

siRNAs targeting *HIF1A* (sc-35561) or *EPAS1* (sc-35316) and a control siRNA (sc-37007) were purchased from Santa Cruz Biotechnology. Lipofectamine RNAiMAX (ThermoFisher: 13778030) was used according to manufacture recommendations. Cells were transfected 24 hr prior to exposure to hypoxic conditions.

### Generation of cells with stable shRNA-mediated knockdowns

Five non-overlapping shRNA sequences that target *NSD2* were chosen from The RNAi Consortium (TRC) shRNA Library (Broad Institute). Forward and reverse oligonucleotides (ThermoFisher: #1033602) for each shRNA insert were annealed for 4 min at 95°C. Annealed oligonucleotides were inserted into a pLKO.1-TRC vector with puromycin resistance gene (*92*) using EcoRI (NEB: #R0101T) and AgeI (NEB: #R3552S) restriction sites. shRNA inserts were ligated into a digested pLKO.1 vector using NEB T4 DNA ligase (NEB: #M0202T) following manufacturer recommendations. Ligated vectors were transformed into competent DH5α cells by heat shocking bacteria in a 42°C water bath for 45 sec. Ampicillin-resistant clones were screened by EcoRI (NEB: #R0101T) and NcoI (NEB: #R0193T) restriction digest and run on a 1% agarose gel. A scrambled control in a pLKO.1 vector was purchased from Sigma-Aldrich (Sigma: #SHC016). 0082T CAFs were transduced with *NSD2*-targeting or control shRNAs using second-generation lentiviral transfection of LX293T cells (Takara: #632180) with pCMV-VSVG (Addgene: #8454) and pCMV-delta8.2 (Addgene: #8455) packaging plasmids. Viral supernatants were passed through 0.45-µm filters and supplemented with 8 μg/mL polybrene prior to use for 0082T cell transduction. Transductants were selected using puromycin (2 μg/mL) for 5 days before screening by immunofluorescence microscopy for loss of NSD2 or H3K36me2 signal. The two shRNAs that produced the greatest knockdown of NSD2 and decreased H3K36me2 (TRCN0000019815 and TRCN0000274232) were used for further experiments.

Four non-overlapping shRNAs targeting *RELA*, which encodes the NF-κB p65 regulatory subunit, were designed using the TRC shRNA library. Forward and reverse sequence oligos for each shRNA insert (ThermoFisher: #1033602) were annealed as previously described. Annealed *RELA* oligos and a control harpin targeting GFP were inserted into the tet-pLKO-puro vector (Addgene: #21915) using EcoRI and AgeI restriction sites. Colony screening for inserts, lentiviral transduction, and 0082T CAF antibiotic selection were conducted as for the *NSD2*-targeting shRNAs. Transduced 0082Ts were treated with 500 ng/mL of doxycycline hyclate (Sigma: #D5207) for 48 hr before assessing NF-κB (p65) knockdown efficiency by immunofluorescence microscopy. The two shRNAs that most effectively depleted NF-κB (TRCN0000014684 and TRCN0000329800) were used for further experiments. Sequences for all *NSD2* and *RELA* shRNAs used in this study are provided in **Supplementary Table S7**.

### Phospho-kinase array

HPAF-II cell lysates were prepared according to manufacturer instructions and analyzed using the Proteome Profiler Human Phospho-Kinase Array (R&D Systems: #ARY003C). The array was developed on X-ray film at multiple exposures (30 sec, 1 min, 3 min, 5 min, 7.5 min, and 10 min) and imaged using a GS-800 Densitometer (Bio-Rad). A 5-min exposure was used for quantification after confirming it fell within the dynamic range by plotting exposure time against optical density for the positive control spots.

### ELISA

Conditioned medium from 0082T cells, prepared and collected as described above, was concentrated using 3-kDa centrifuge filters. Samples were sonicated for 30 sec to disrupt any exosomes before analysis with ELISA kits for HGF (RayBiotech: ELH-HGF-1) or IGF-2 (RayBiotech: ELH-IGF2-1). Sample absorbances were read on a Cytation5 plate reader (BioTek), and absolute concentrations were computed by via calibration curves generated using recombinant proteins.

### Single-cell RNA sequencing data analysis

Annotated and pre-processed PDAC scRNA-seq data acquired by Elyada et al. (*41*) were kindly provided by Dr. David Tuveson (Cold Spring Harbor Laboratory). Ductal cell clusters 1 (n=902), 2 (n=264), and 3 (n=92) identified by those authors were combined to make one cancer cell population for our analysis using the *Seurat* R package (v5.0.2) (*93*). All ductal cell clusters were aggregated with the CAF-enriched fraction, which contained iCAF (n=191 cells) and myCAF (n=789 cells) clusters before additional data analysis. scRNA-seq data were analyzed by gene set variation analysis (GSVA) using the *GSVA* (v1.40.1) R (v4.2.2) package to determine gene set enrichment scores within the CAF population.

### CellPhoneDB, NicheNet, and CellChat analysis

The combined dataset containing malignant cell and CAF clusters was exported as a text file containing count data and cell-type annotations and analyzed using the *CellPhoneDB* (v2.1.0) Python (v2.3.5) package (*52*) or the *CellChat* (v1.5) R package. Growth factors and cytokines were designated as iCAF- or myCAF-secreted based on the CAF subtype that exhibited greater expression. Interactions identified to occur between iCAFs/myCAFs and ductal cells (cancer cells) from CellPhoneDB were used for downstream NicheNet analysis (*54*). The NicheNet database (v1.1.1) for predicted regulatory potential of growth factors and cytokines on genes was used to generate scores for each growth factor or cytokine for patient-derived mesenchymal marker genes (*55*) and NF-κB target genes (*94*). Scores for mesenchymal genes for each CAF-expressed cytokine or growth factor were summed to generate an integrated score. Integrated scores were ranked to identify the cytokines or growth factors predicted to contribute to CAF-mediated EMT. Integrated scores for NF-κB target genes were similarly determined for ductal cell-secreted factors predicted to promote signaling in CAFs.

### scRNA-seq clustering

Clustering and low-dimensional projections were used as two methods to simplify high-dimensional scRNAseq data (*95*). Hierarchical clustering using Ward’s method (Ward.D2 algorithm) was performed on the PDAC CAFs using only gene expression for cytokines and growth factors compiled by the Molecular Signatures database (*94*) and plotted as a heatmap using the *pheatmap* (v1.0.12) R package. A 2D UMAP projection (nearest neighbors setting of 15 and minimum distance setting of 0.01) was created for CAF scRNA-seq data using Hallmark Hypoxia genes as features (*47*). Consensus clustering (*ConsensusClusterPlus* R package; v3.18) was then used to identify *k* = 2 as the optimal number of clusters, which were classified as hypoxia-high (n=176) or -low (n=777) based on Hallmark Hypoxia UCell scores (*Ucell* R package; v2.6) (*96*). Consensus clustering was performed using Euclidean distance, Ward’s linkage for subsampling, and the partitioning around medoids algorithm.

### Statistical analyses

All statistical analyses were performed using R version 4.2.2. For all experiments where statistics were computed, at least three biological replicates were analyzed. An unpaired two-tailed Student’s t-test was performed for qRT-PCR and cell morphology analyses involving one comparison. A paired two-tailed Student’s t-test was used to compare protein expression in Hypoxyprobe-positive and -negative CAFs within the same KPCY tumor sample. A Wilcoxon rank-sum test was performed for scRNA-seq analyses comparing gene expression between two cell populations. For experiments comparing signal-positive fractions of cells across three or more conditions, a one-way ANOVA with Tukey’s multiple comparison post-hoc test was performed. For cell morphology studies involving single-cell measurements across three or more conditions, a one-way ANOVA with Games-Howell post-hoc test was used.

## ACKNOWLEDGEMENTS

We thank Dr. John Lazo and Dr. Elizabeth Sharlow for providing the kinase inhibitor library. We thank Dr. Andrey Lowy for generating the 0082T immortalized CAF cell line and Dr. Mara Sherman for sharing it. We acknowledge Dr. David Tuveson (Cold Springs Harbor Laboratory) for kindly sharing the annotated scRNA-seq data set used in our analysis. We thank Paul Myers for technical discussions regarding scRNA-seq data analysis. We thank Logan Campbell for technical discussion on designing shRNA plasmids. **Figure 2E, 3C** and **Figure 7** were created using BioRender.com.

## AUTHOR CONTRIBUTIONS

**K.M. Kowalewski**: Conceptualization, investigation, methodology, data analysis, writing-original draft, and editing. **S.J. Adair**: Investigation, methodology, writing-review and editing. **A. Talkington**: Methodology, writing-review and editing. **J.J. Wieder**: Investigation and methodology. **J.R. Pitarresi**: Investigation, methodology, writing-review and editing. **K. Perez-Vale**: Investigation, methodology. **B. Chu**: Investigation and methodology. **S. Dolatshahi**: Conceptualization and supervision. **R. Sears**: Resources, writing-review and editing. **B.Z. Stanger**: Conceptualization, resources, supervision, writing-review and editing. **T.W. Bauer**: Conceptualization, resources, supervision, writing-review and editing. **M.J. Lazzara**: Conceptualization, supervision, funding-acquisition, project administration, writing-original draft, writing-review and editing.

## MATERIALS & CORRESPONDENCE

Correspondence to Matthew J. Lazzara.

## SUPPLEMENTARY FIGURES

**Supplementary Figure S1.**
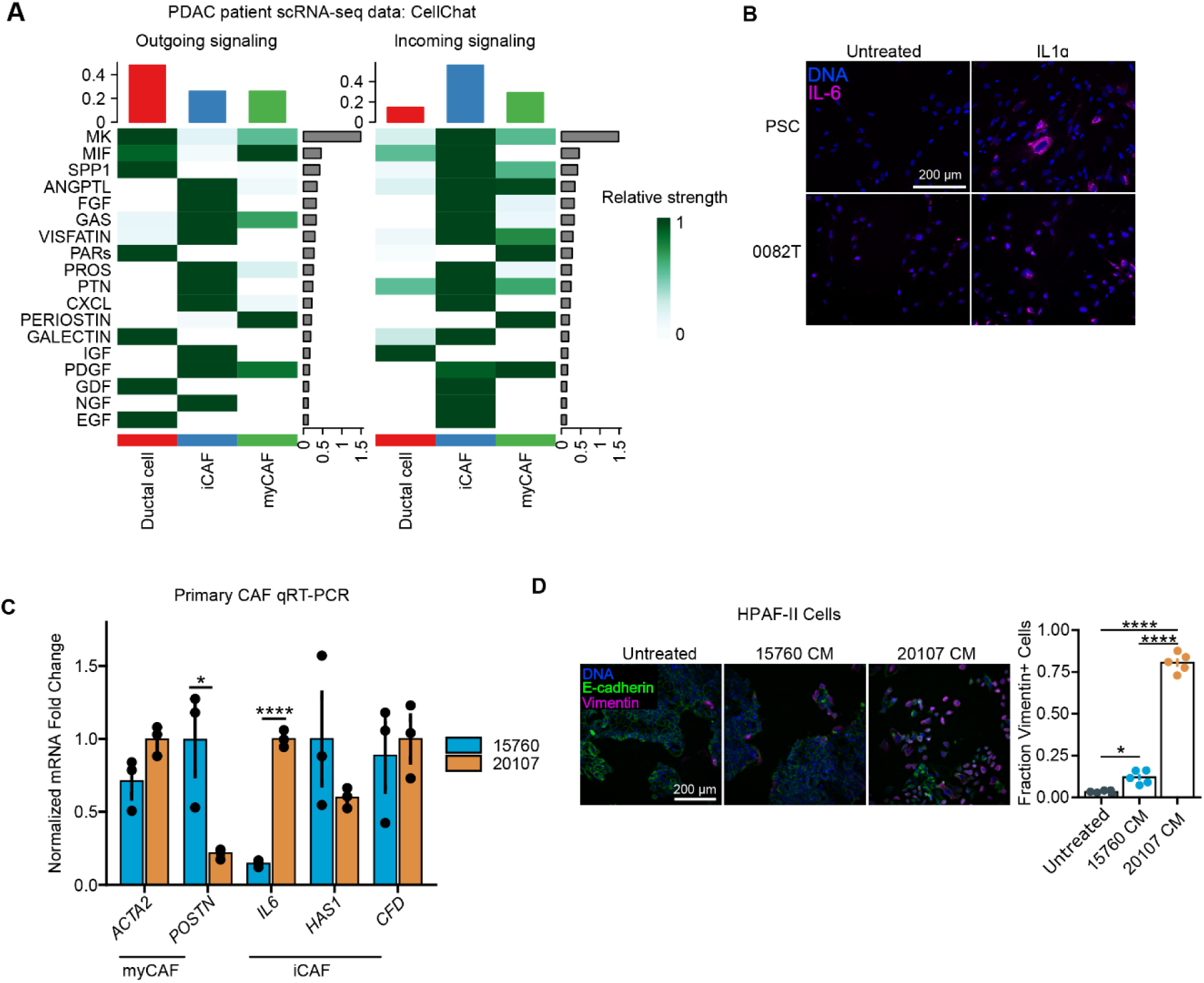
iCAFs express numerous growth factors/cytokines and preferentially promote EMT in cancer cells. **A)** CellChat analysis was performed using previously acquired scRNA-seq (*41*) data to predict growth factor/cytokine signaling interactions between iCAFs-myCAFs and ductal cells (cancer cells). **B)** Immunofluorescence microscopy for IL-6 was performed on PSCs and 0082T CAFs that were treated with or without IL1α (10 ng/mL) for 72 hr. Images representative of n=3. **C)** qRT-PCR for the indicated transcripts was performed on mRNA isolated from primary human CAF lines 20107 and 15760. n=3, t-test. **D)** HPAF-II cells were treated with conditioned medium (CM) that was generated from 20107 cells (iCAF-like) or 15760 cells (myCAF-like) for 120 hr, and immunofluorescence microscopy was performed for the indicated proteins. n=3, one-way ANOVA with Tukey’s multiple comparison test. Data are presented as mean ± SEM (C, D). * *p* <0.05, ** *p* < 0.01, *** *p* < 0.001, **** *p* < 0.0001

**Supplementary Figure S2.**
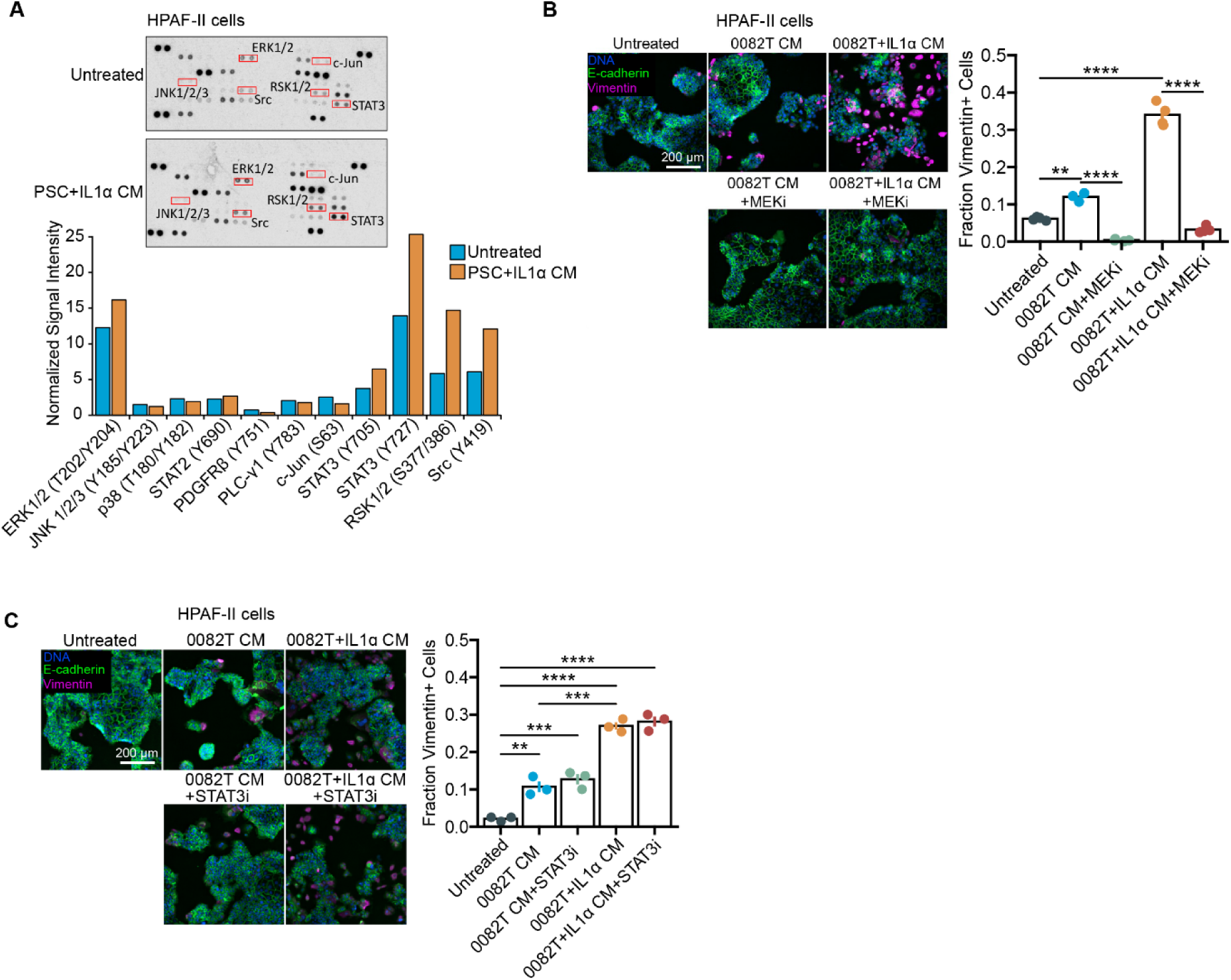
ERK activation by CAF-conditioned medium promotes EMT in cancer cells. **A)** HPAF-II cells were treated with PSC+IL1α conditioned medium (CM) or untreated for 24 hr, and lysates were analyzed using a Proteome Profiler Phospho-kinase Array. Image for n = 1. **B)** HPAF-II cells were treated with CM from 0082T cells treated with or without IL1α in combination with DMSO or CI-1040 (MEKi) for 120 hr. Immunofluorescence microscopy was performed for the indicated proteins. n=3, one-way ANOVA with Tukey’s multiple comparison test. **C)** HPAF-II cells were treated with CM from 0082T cells treated with or without IL1α in combination with DMSO or STATTIC (STAT3i, 1 μM) for 120 hr. Immunofluorescence microscopy was performed for the indicated proteins. n=3, one-way ANOVA with Tukey’s multiple comparison test. Data presented as mean ± SEM (B, C). ** *p* < 0.01, *** *p* < 0.001, **** *p* < 0.0001

**Supplementary Figure S3.**
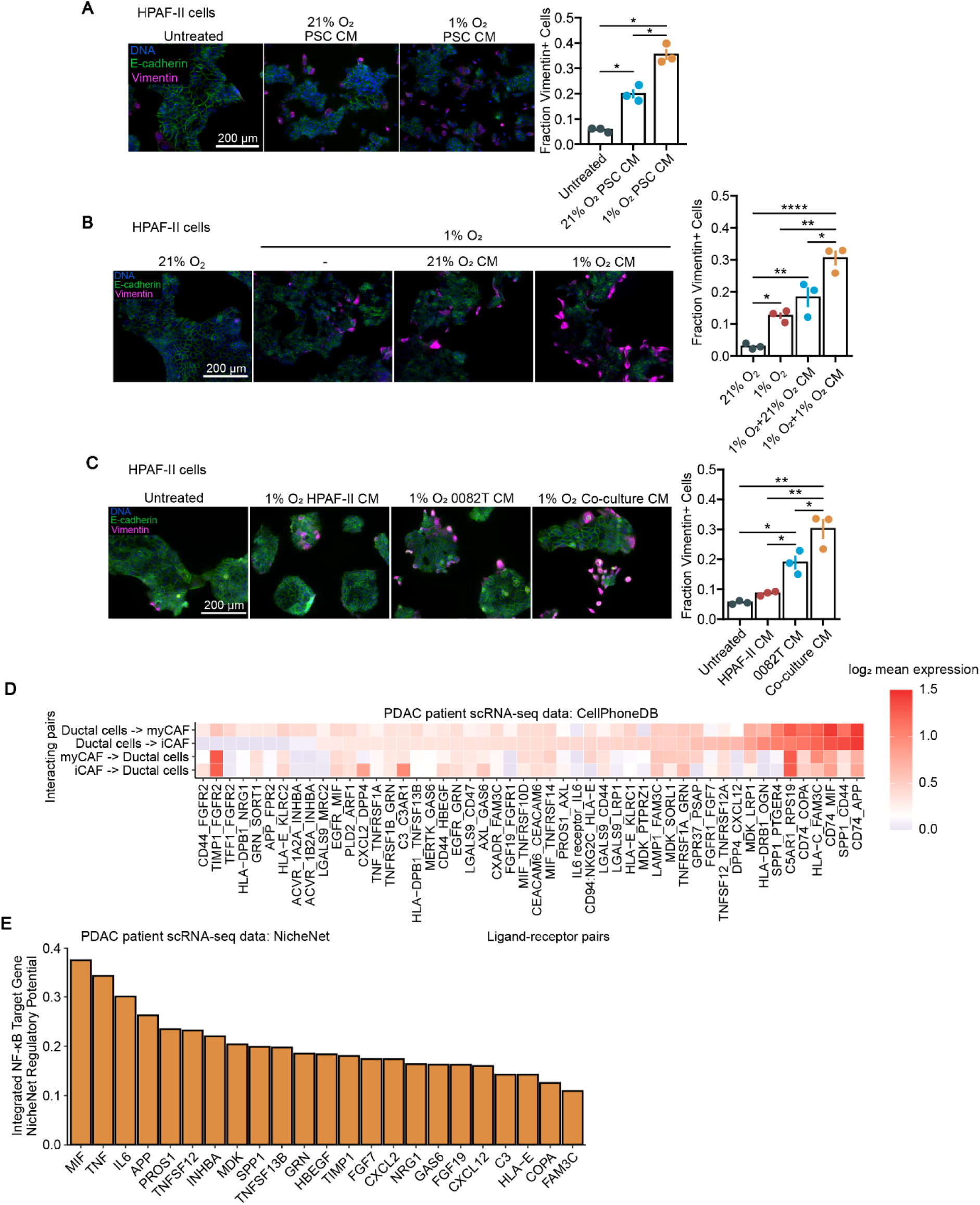
Hypoxia enhances the ability of CAFs to promote EMT and cooperates with hypoxia-driven cell-autonomous EMT in cancer cells. **A)** HPAF-II cells were treated with conditioned medium (CM) from human PSCs for 120 hr, and immunofluorescence microscopy for the indicated proteins was performed. n=3, one-way ANOVA with Tukey’s multiple comparison test. **B)** HPAF-II cells were cultured in 21% or 1% O_2_ for 120 hr. For the 1% O_2_ condition, HPAF-II cells were treated with CM from 0082T cells that had been cultured at 21% or 1% O_2_ for 120 hr or were untreated. n=3, one-way ANOVA with Tukey’s multiple comparison test. **C)** HPAF-II cells were treated for 120 hr with CM generated from HPAF-II cells, 0082T cells, or a co-culture of HPAF-II and 0082T cells. n=3, ANOVA with Tukey’s multiple comparison test. **D)** Using the scRNA-seq data analyzed in Figure 1 (*41*), ligand-receptor interactions between ductal cells and CAFs were predicted using CellPhoneDB analysis and filtered for the top 50 ductal/CAF cell interactions. **E)** Growth factor and cytokine regulatory potential for Hallmark NF-κB target genes compiled by the Molecular Signatures Database (*94*) was determined for ligands nominated by prior CellPhoneDB analysis in (D). Integrated scores for each ligand were calculated by summing the NicheNet regulatory potential for each gene contained in the Hallmark NF-κB gene set. Data are presented as mean ± SEM (A, B, C). * *p* <0.05, ** *p* < 0.01, *** *p* < 0.001, **** *p* < 0.0001

**Supplementary Figure S4.**
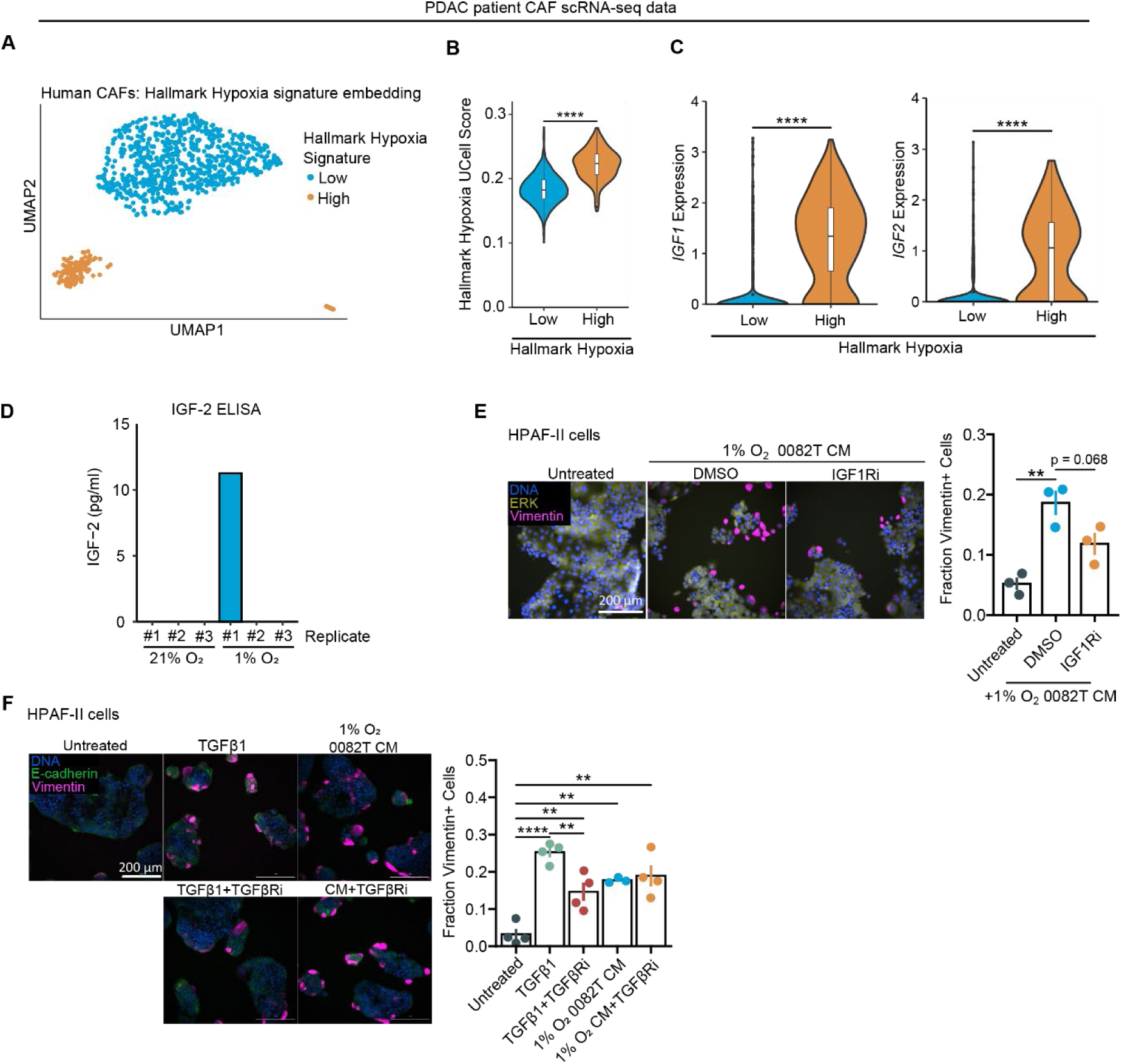
Hypoxic CAFs support cancer cell EMT through IGF-2 secretion. **A)** Using previously acquired scRNA-seq data (*41*), consensus clustering was performed on CAFs using only Hallmark Hypoxia genes and visualized by UMAP projection. **B)** UCell scores were used to quantify enrichment of Hallmark Hypoxia genes for cells in clusters identified in panel (A). Wilcoxon rank-sum test. **C)** For the clusters described in panel (A), *IGF1* and *IGF2* expression were assessed. Wilcoxon rank-sum test. **D)** Conditioned medium was collected from 0082T cells cultured in 21% or 1% O_2_ and analyzed for IGF-2 by ELISA. n=3. **E)** HPAF-II cells were treated for 120 hr with conditioned medium (CM) from hypoxic 0082T cells treated with DMSO or linsitinib (2.5 μM). Immunofluorescence microscopy for the indicated proteins was performed. n=3, one-way ANOVA with Tukey’s multiple comparison test. **F)** HPAF-II cells were treated for 120 hr with TGFβ1 (10 ng/mL) or CM from hypoxic 0082T cells ± galunisertib (TGFβRi; 10 μM). Immunofluorescence microscopy was performed for the indicated proteins. n=3, one-way ANOVA with Tukey’s multiple comparison test. Data are presented as mean ± SEM for bar plots (E, F), as violin plots overlayed with box-and-whisker plots that represent the first quartile (lower box bound), median (center bound), or third quartile (upper box bound) (B, C) * *p* <0.05, ** *p* < 0.01, *** *p* < 0.001, **** *p* < 0.0001

**Supplementary Figure S5.**
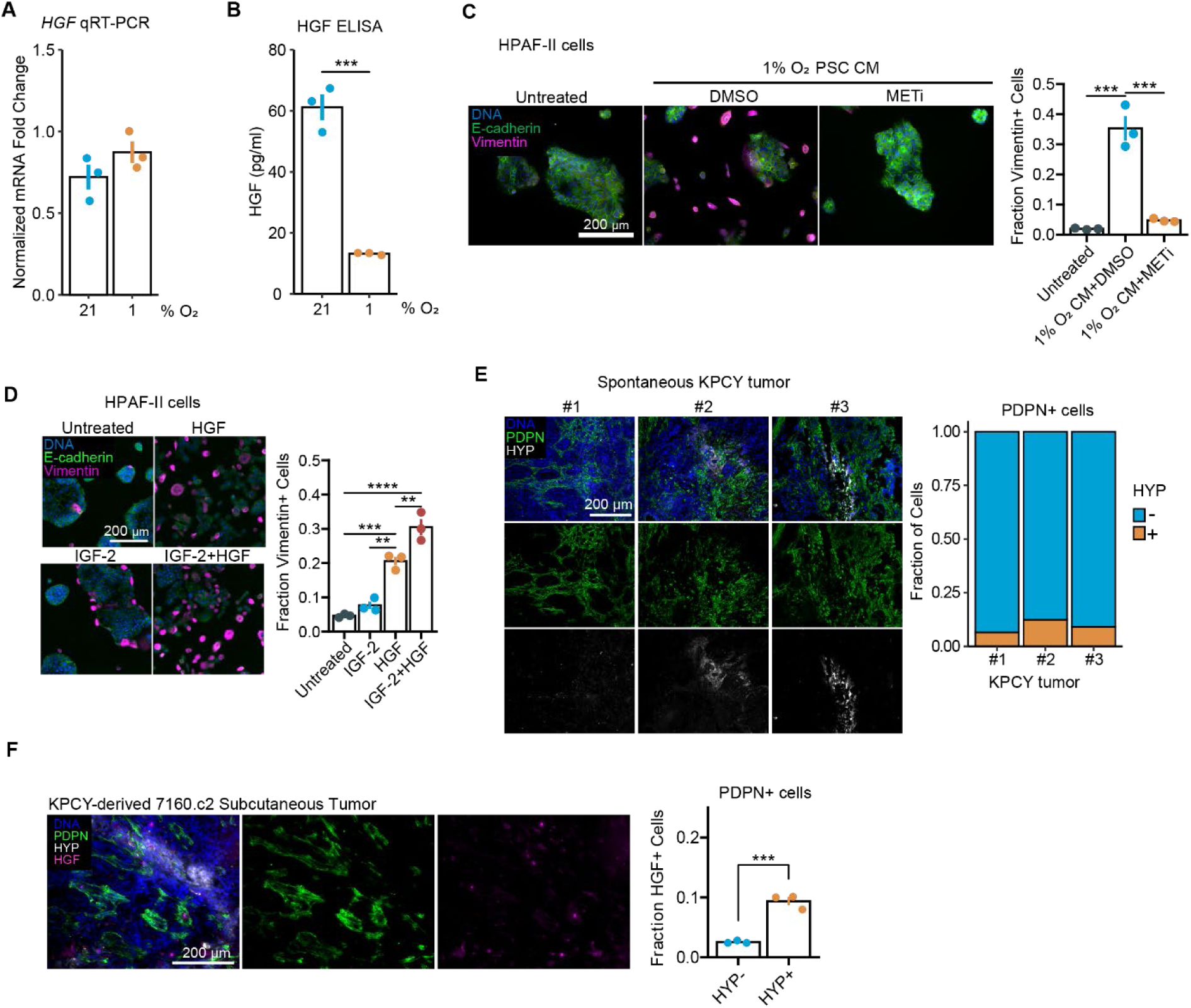
HGF secreted by hypoxic CAFs facilitates cancer cell EMT. **A)** 0082T CAFs were cultured in 21% or 1% O_2_ for 120 hr, and qRT-PCR was performed on extracted mRNA for *HGF* transcripts. *CASC3* was used for normalization. n=3. **B)** Conditioned medium was collected from 0082T cells cultured in 21% or 1% O_2_ and analyzed for HGF by ELISA. n=3, t-test. **C)** HPAF-II cells were untreated or treated with conditioned medium (CM) from PSCs grown at 1% O_2_ +/- PHA665752 (METi, 2.5 μM) for 120 hr. Immunofluorescence microscopy was performed for the indicated proteins. n=3, one-way ANOVA with Tukey’s multiple comparison test. **D)** HPAF-II cells were treated with IGF-2 (100 ng/mL), HGF (5 ng/mL), or IGF-2+HGF for 120 hr. Immunofluorescence microscopy was performed for the indicated proteins. n=3, one-way ANOVA with Tukey’s multiple comparison test. **E)** Fluorescence-based immunohistochemistry was performed on sections of KPCY tumors for podoplanin (PDPN) and Hypoxyprobe (HYP) to calculate the percentage of CAFs within hypoxic regions. n=3. **F)** In sections of subcutaneous tumors formed using KPCY-derived 7160.c2 cells, HGF expression was assessed in Hypoxyprobe-positive and -negative CAFs (PDPN+ cells). n=3, t-test. Data are presented as mean ± SEM for bar plots (A, B, C, D, F) or as a mosaic plot indicating the percentage of PDPN+ cells that are positive or negative for Hypoxyprobe (HYP) across biological replicates. * *p* <0.05, ** *p* < 0.01, *** *p* < 0.001, **** *p* < 0.0001

**Supplementary Figure S6.**
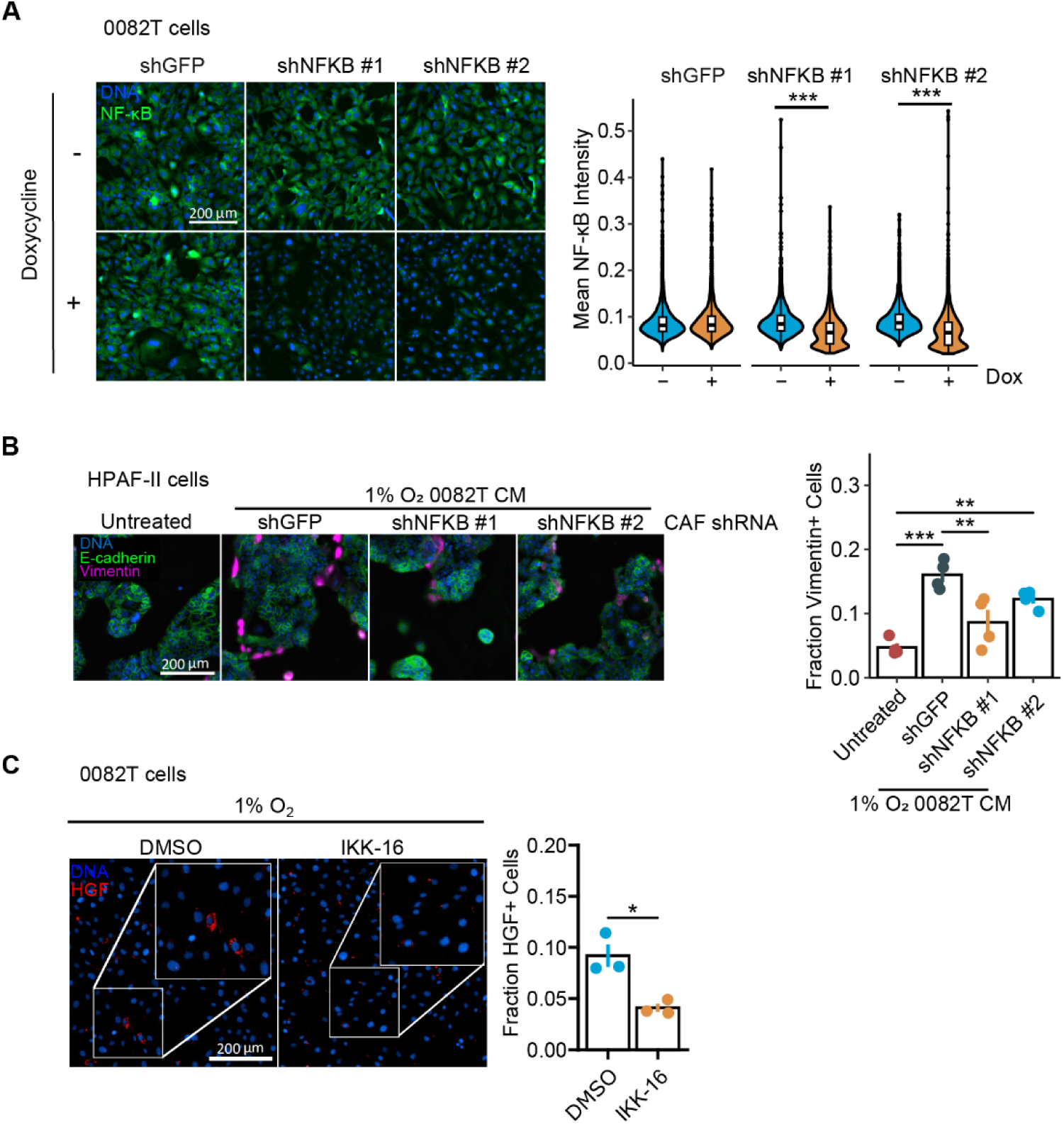
NF-κB promotes HGF expression in hypoxic CAFs and supports the ability of hypoxic CAFs to promote EMT. **A)** 0082T CAFs engineered with doxycycline-inducible shRNAs for knockdown of NF-κB (p65) or GFP were treated with 500 ng/mL doxycycline for 48 hr. Immunofluorescence microscopy was performed for NF-κB (p65). n=3, t-test. **B)** HPAF-II cells were untreated or treated for 120 hr with conditioned medium (CM) from the 0082T cells described in (A) that were grown at 1% O_2_ and doxycycline-treated for 48 hr. Immunofluorescence microscopy was performed for the indicated proteins. n=3, one-way ANOVA with Tukey’s multiple comparison test. **C)** 0082T CAFs were cultured in 1% O_2_ and treated with DMSO or IKK-16 (0.5 μM) for 120 hr. n=3, t-test. Data are presented as mean ± SEM for bar plots (B, C), as violin plots overlayed with box-and-whisker plots that represent the first quartile (lower box bound), median (center bound), or third quartile (upper box bound) (A), * *p* <0.05, ** *p* < 0.01, *** *p* < 0.001

**Supplementary Figure S7.**
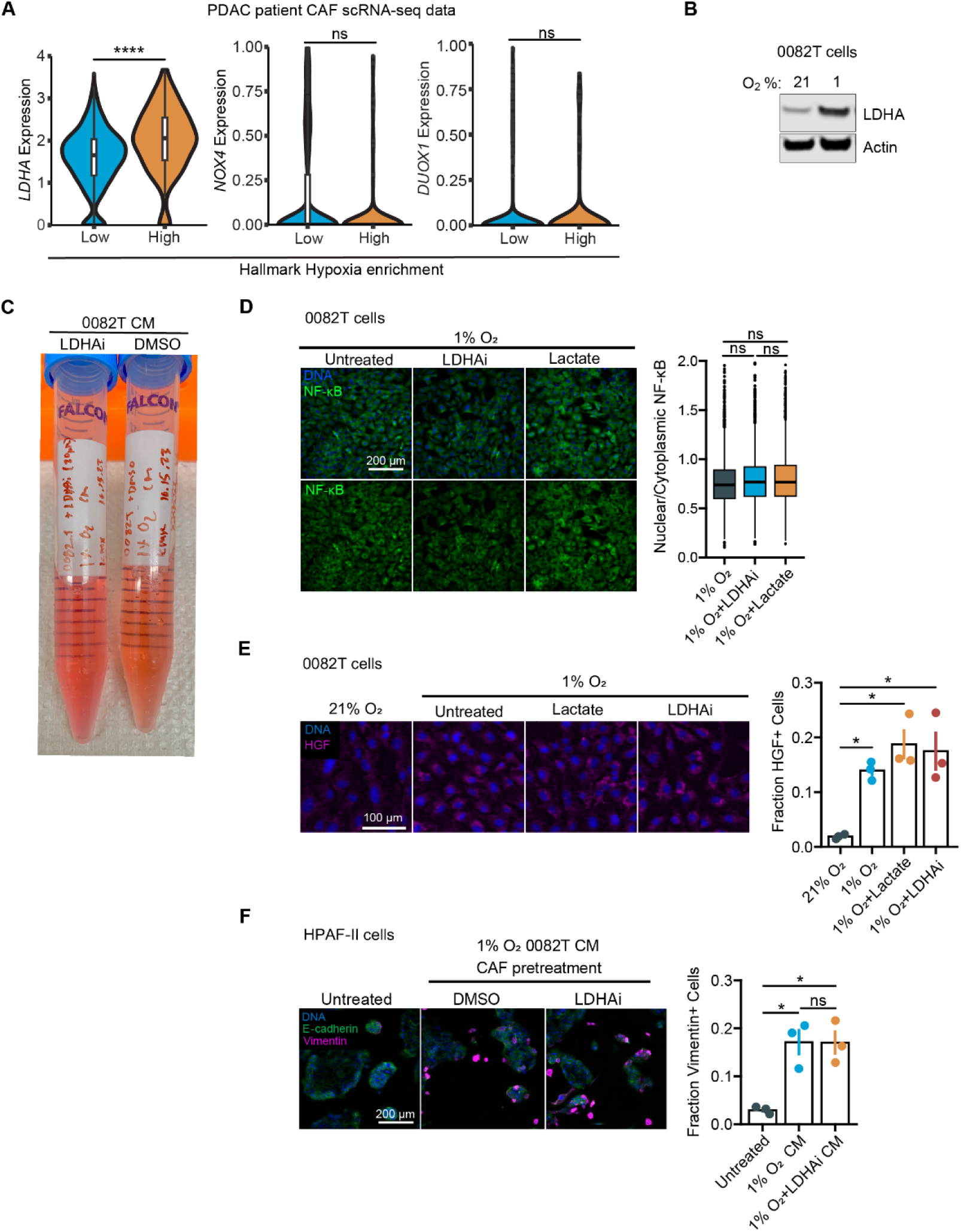
LDHA does not regulate NF-κB in hypoxic CAFs or enhance the ability of CAFs to promote EMT. **A)** Using previously acquired scRNA-seq (*41*) CAFs were classified for low or high enrichment of Hallmark Hypoxia genes and then assessed for expression of the indicated reactive oxygen species-producing enzymes. Wilcoxon rank-sum test. **B)** 0082T CAFs were cultured in 21% or 1% O_2_ for 120 hr, and Western blotting was performed on lysates for the indicated proteins. Image representative of n = 3. **C)** 0082T CAFs were treated with DMSO or GSK2837808A (LDHAi, 10 μM) for 72 hr and used to generate conditioned medium (CM). pH was qualitatively indicated by the color of the phenol-red containing medium (lighter color, indicating lower pH). **D)** 0082T cells were cultured in 1% O_2_ and treated with GSK2837808A (LDHAi, 10 μM) or lactate (30 mM) for 120 hr. Immunofluorescence microscopy was performed for NF-κB to assess nuclear/cytoplasmic ratio. n=3, one-way ANOVA with Tukey’s multiple comparison test. **E)** 0082T CAFs cultured in 21% O_2_ or 1% O_2_ and were treated with DMSO, lactate (30 mM), or GSK2837808A (LDHAi, 10 μM) for 120 hr. Immunofluorescence microscopy was performed for HGF. n=3, one-way ANOVA with Tukey’s multiple comparison test. **F)** HPAF-II cells were treated for 120 hr with or without CM that was generated from 0082T CAFs pretreated with DMSO or GSK2837808A (LDHAi, 10 μM). n=3, one-way ANOVA with Tukey’s multiple comparison test. Data are presented as mean ± SEM for bar plots (E, F), as violin plots overlayed with box-and-whisker plots that represent the first quartile (lower box bound), median (center bound), or third quartile (upper box bound) (A), or as a box-and-whisker plot only (D). * *p* <0.05, **** p<0.0001

**Supplementary Figure S8.**
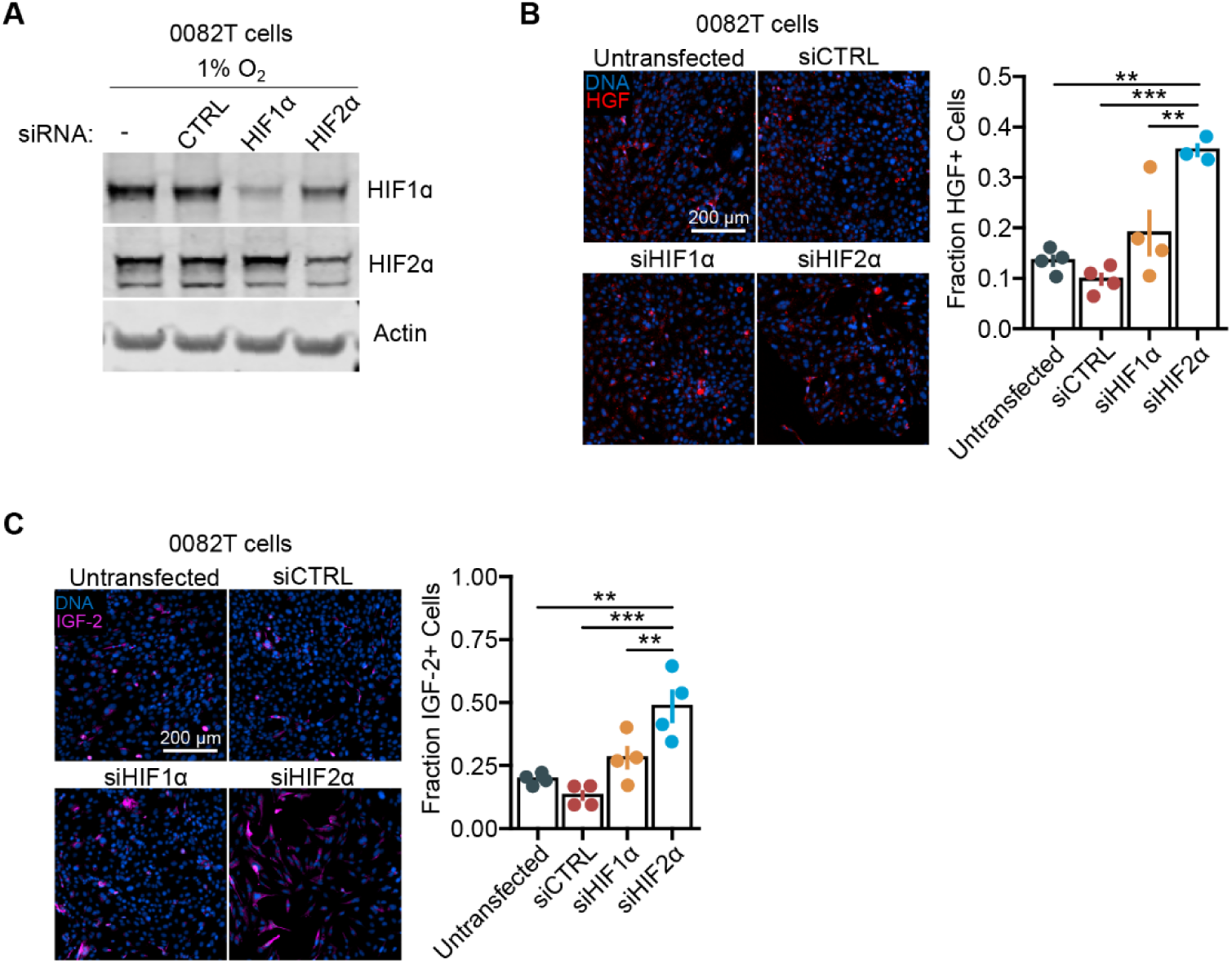
HIF1α and HIF2α do not promote HGF or IGF-2 expression in hypoxic CAFs. **A)** 0082T cells were transfected with siRNA against *HIF1A*, siRNA against *EPAS1* (gene encoding HIF2α), or a control siRNA and cultured in 1% O_2_ for 4 hr before lysing. Western blotting was performed for the indicated proteins. Image representative of n=3. **B)** and **C)** 0082T CAFs were transfected with the siRNAs described in (A) and grown for 120 hr at 1% O_2_. Immunofluorescence for HGF or IGF-2 was performed. Data are presented as mean ± SEM. Throughout the panels, ** *p* < 0.01, *** *p* < 0.001.

**Supplementary Figure S9.**
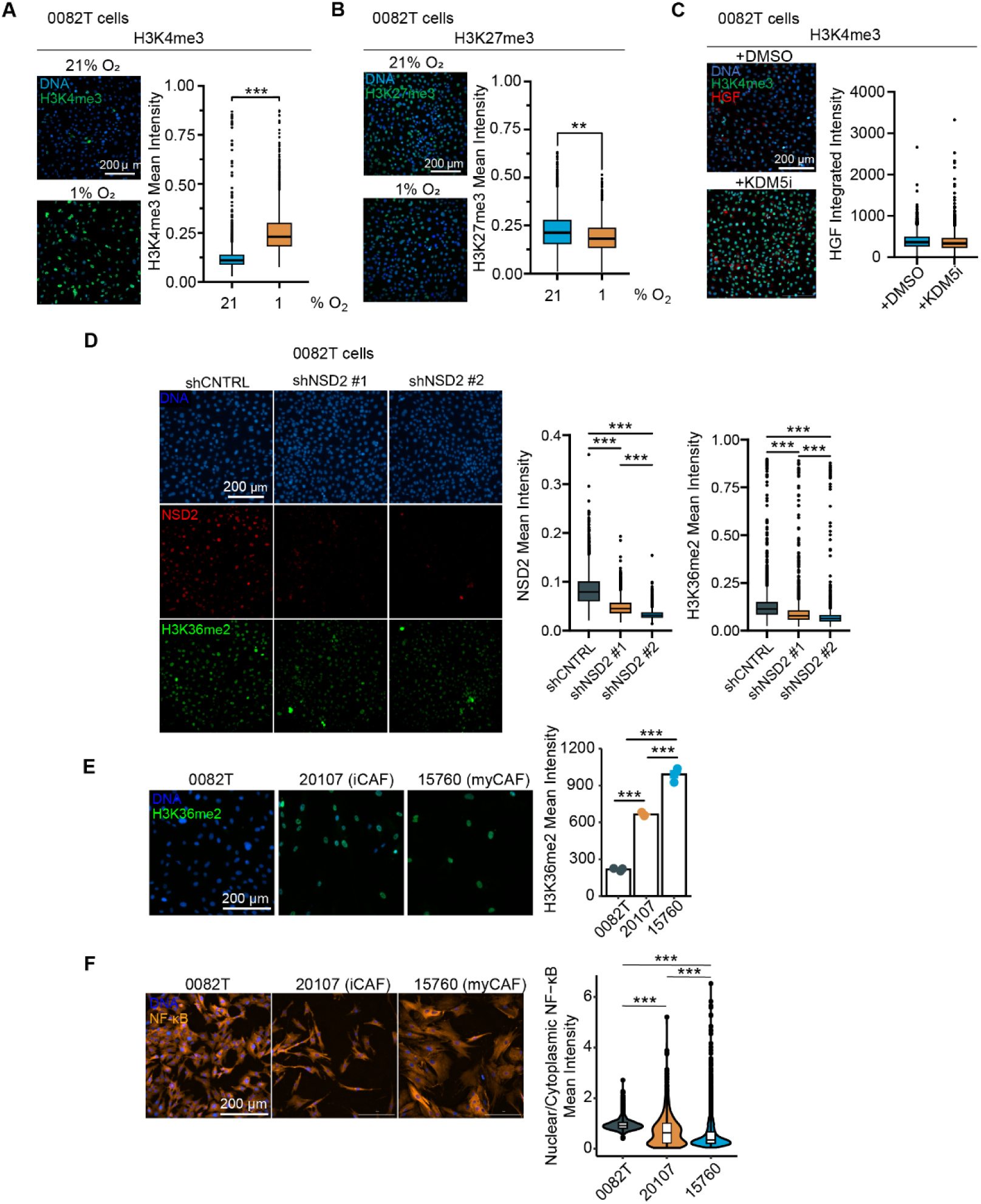
Hypoxia alters histone methylation in CAFs. **A)** and **B)** 0082T CAFs were cultured at 21% or 1% O_2_ for 120 hr, and immunofluorescence microscopy for histone 3 lysine 4 trimethylation (H3K4me3) or histone 3 lysine 27 trimethylation (H3K27me3) was performed. n=3, t-test. **C)** 0082T CAFs were cultured in 1% O_2_ with DMSO or CPI-455 (KDM5i) for 72 hr and analyzed for H3K4me3 and HGF expression by immunofluorescence microscopy. n=3, t-test. **D)** 0082T engineered with shRNAs targeting NSD2 or a scrambled control (shCNTRL) were cultured in 1% O_2_ and stained for NSD2 and H3K36me2 by immunofluorescence microscopy. n=5, one-way ANOVA with Tukey’s multiple comparison test. **E)** and **F)** Two primary human PDAC CAF lines (15760 and 20107) and 0082T CAFs were cultured at 21% O_2_ and analyzed for H3K36me2 and NF-κB (p65) by immunofluorescence microscopy. n=3, one-way ANOVA with Tukey’s multiple comparison test. Data are presented as mean ± SEM for bar plots (E, F), as box-and-whisker plots that represent the first quartile (lower box bound), median (center bound), or third quartile (upper box bound) (A, B, C, D), or as a violin plot overlayed with a box-and-whisker plot (F). ** *p* < 0.01, *** *p* < 0.0001

**Supplementary Figure S10.**
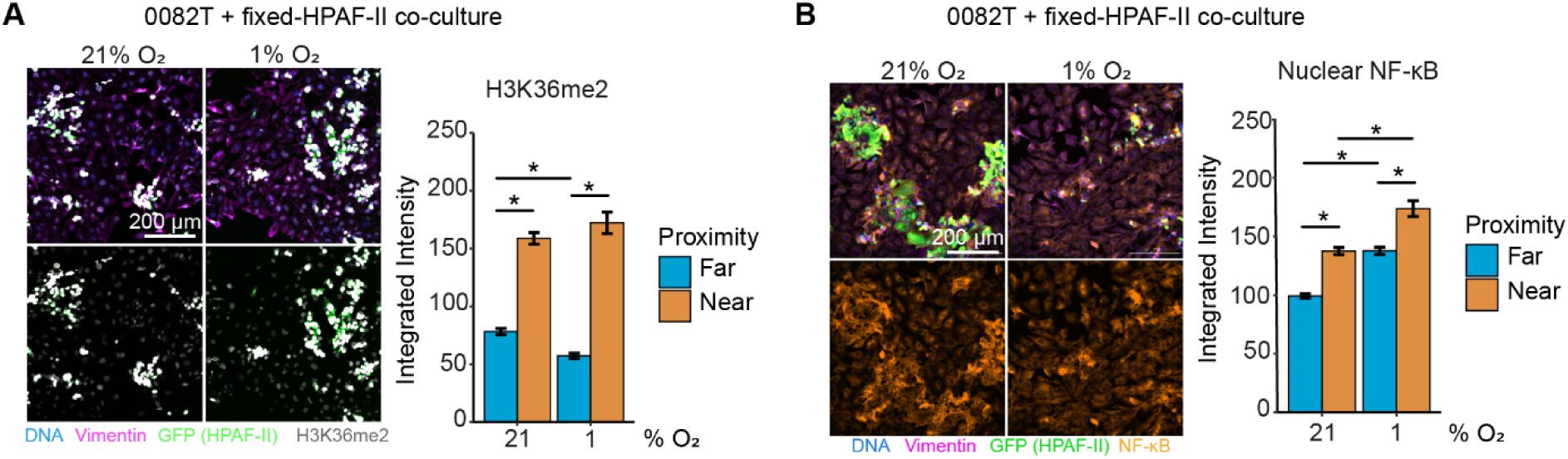
Juxtracrine signaling between HPAF-II cancer cells and 0082T CAFs promotes NF-κB nuclear localization and H3K36 dimethylation in CAFs. 0082T cells were cultured in 21% or 1% O_2_ before co-culturing with PFA-fixed GFP-labelled HPAF-II for 48 hr. **A)** and **B)** Co-cultures were stained for H3K36me2 (A) or NF-κB (B) 48 hr after the addition of 0082T cells, and the resulting images were used to compute distances between 0082T and HPAF-II cells. 0082T cells were binned as being near (<20 μm) or far (>20 μm) from HPAF-II cells. n=3, two-way ANOVA. Data are presented as mean ± standard deviation. Throughout the figure, * *p* < 0.05.

**Supplementary Figure S11.**
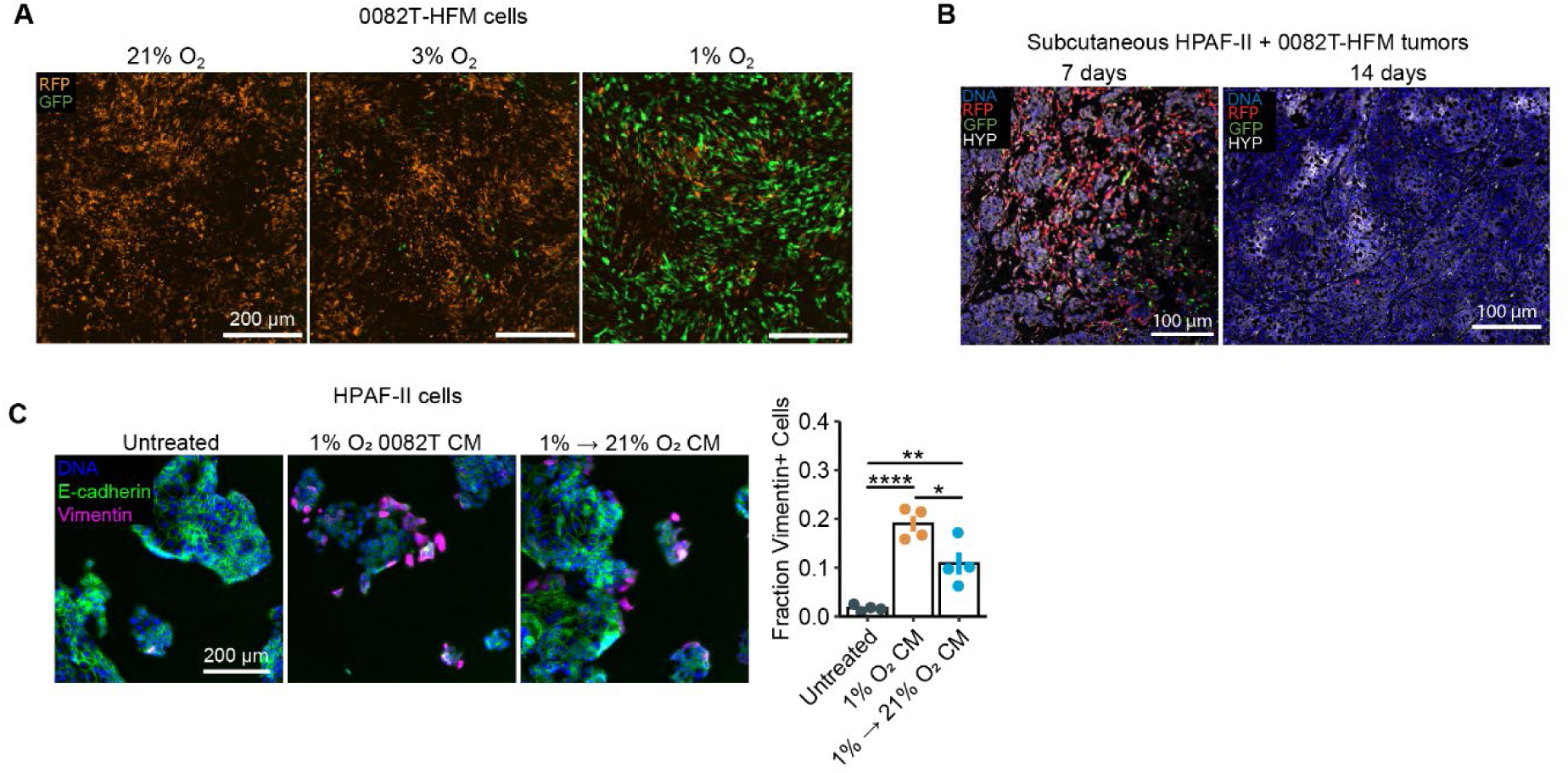
The hypoxia-induced pro-EMT CAF phenotype is reversible. **A)** 0082T-HFM CAFs were cultured at the indicated oxygen concentrations for 7 days and analyzed for GFP and DsRed expression by live-cell imaging. Images representative of n=3. **B)** HPAF-II and 0082T-HFM cells were used to form subcutaneous tumors in mice. Tumors were harvested 7 or 14 days post-implantation. Fluorescence-based immunohistochemistry was performed on tumor sections for the indicated proteins and for Hypoxyprobe (HYP). n=4. **C)** HPAF-II cells were treated with conditioned medium (CM) from 0082T CAFs cultured in 1% O_2_ or 0082T CAFs that were reoxygenated (1% → 21% O_2_) and analyzed by immunofluorescence microscopy for the indicated proteins. n=4, one-way ANOVA with Tukey’s multiple comparison test. Data are presented as mean ± SEM for bar plot (C) or as violin plots overlayed with box- and-whisker plots that represent the first quartile (lower box bound), median (center bound), or third quartile (upper box bound) * *p* < 0.05, ** *p* < 0.01, **** *p* < 0.0001

## SUPPLEMENTARY TABLES

**Supplementary Table S1.**
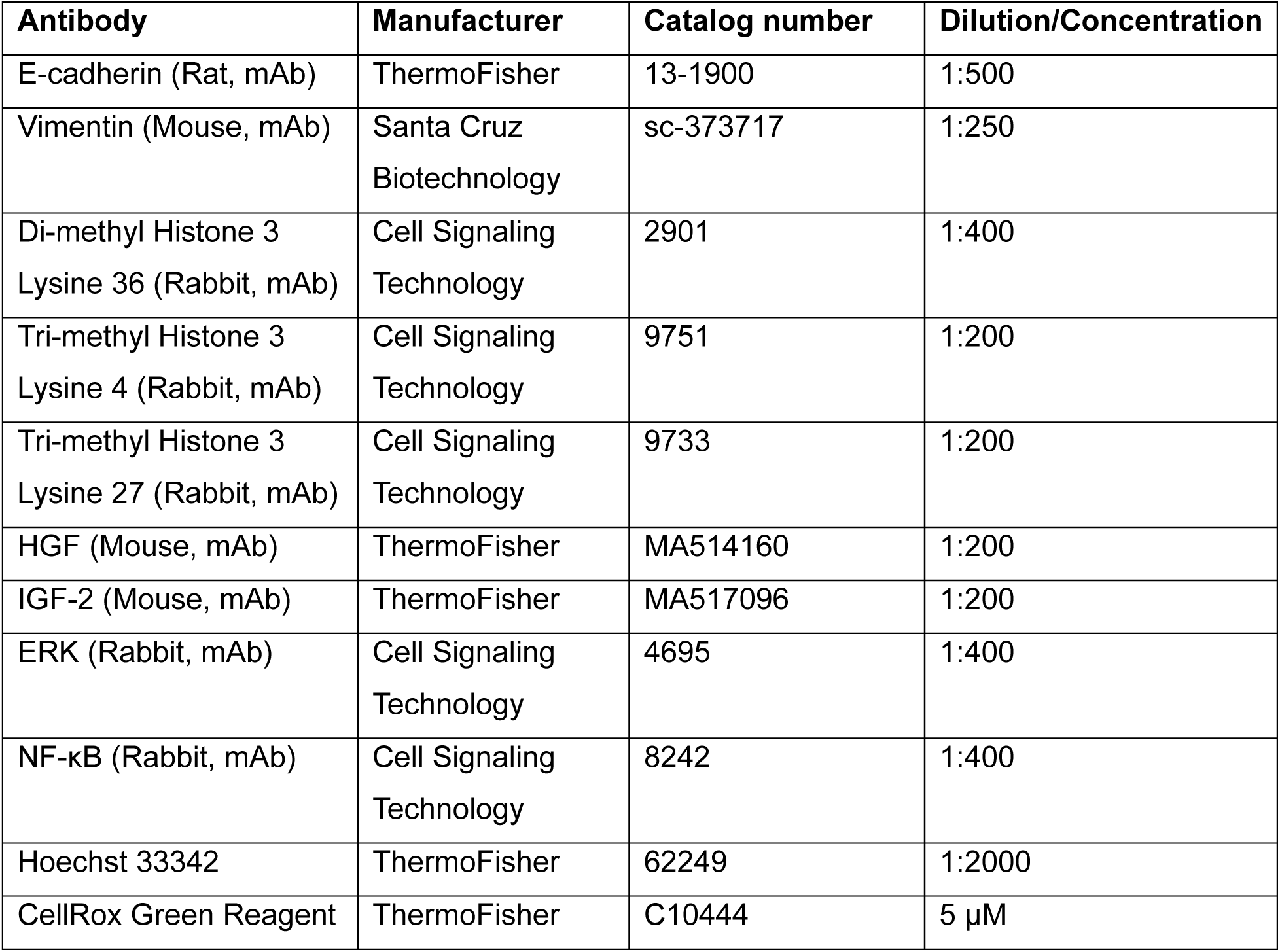
Antibodies and dyes used for immunofluorescence microscopy.

**Supplementary Table S2.**
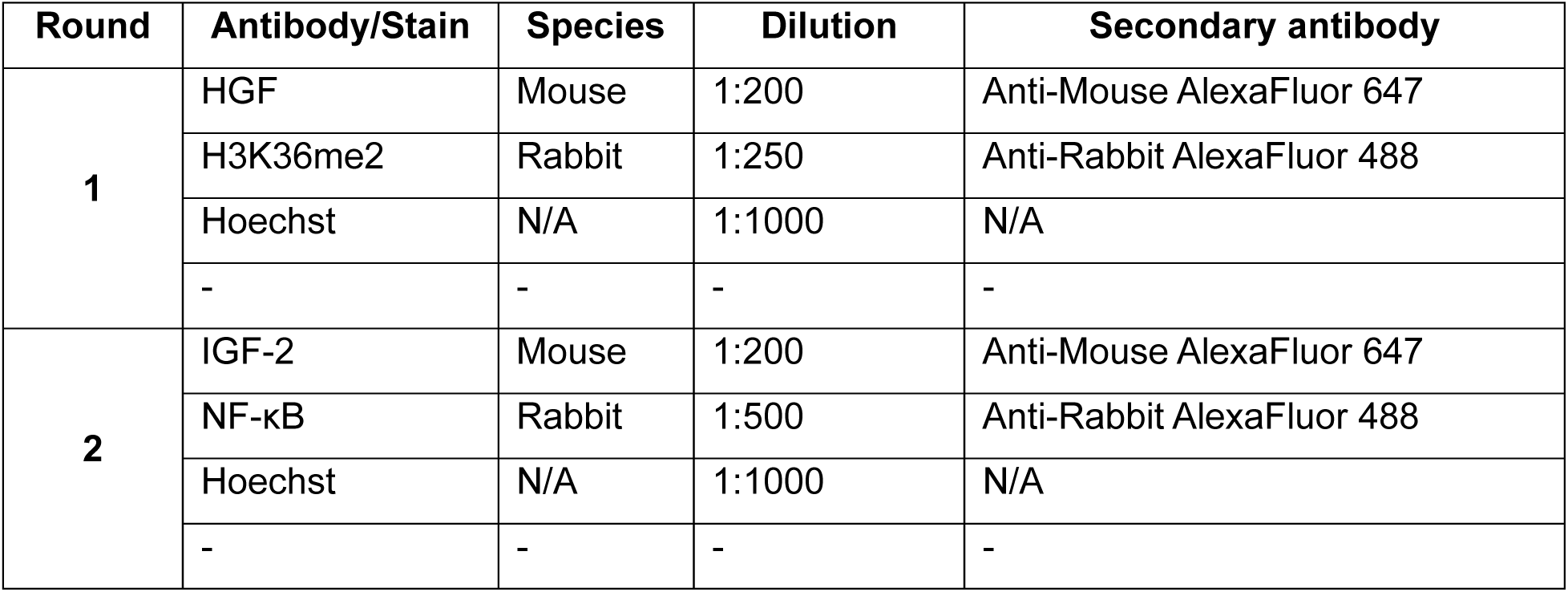
4i staining for 0082T HGF, IGF-2, NF-κB, and H3K36me2.

| Round | Antibody/Stain | Species | Dilution | Secondary antibody |
| --- | --- | --- | --- | --- |
| **1** | HGF | Mouse | 1:200 | Anti-Mouse AlexaFluor 647 |
|  | H3K36me2 | Rabbit | 1:250 | Anti-Rabbit AlexaFluor 488 |
|  | Hoechst | N/A | 1:1000 | N/A |
|  | - | - | - | - |
| **2** | IGF-2 | Mouse | 1:200 | Anti-Mouse AlexaFluor 647 |
|  | NF-κB | Rabbit | 1:500 | Anti-Rabbit AlexaFluor 488 |
|  | Hoechst | N/A | 1:1000 | N/A |
|  | - | - | - | - |

**Supplementary Table S3.** 4i staining for KPCY tumors for measuring CAF H3K36me2 and NF-κB.

| Round | Antibody/Stain | Species | Dilution | Secondary antibody |
| --- | --- | --- | --- | --- |
| **1** | Podoplanin | Syrian Hamster | 1:200 | Anti-Hamster AlexaFluor 488 |
|  | Hypoxypore | Rabbit | 1:100 | Anti-Rabbit AlexaFluor 546 |
|  | HGF | Mouse | 1:100 | Anti-Mouse 647 |
|  | Hoechst | N/A | 1:1000 | N/A |
| **2** | NF-κB | Rabbit | 1:200 | Anti-Rabbit AlexaFluor 546 |
|  | GFP | Goat | 1:500 | Anti-Goat AlexaFluor 647 |
|  | Hoechst | N/A | 1:1000 | N/A |
|  | - | - | - | - |
| **3** | H3K36me2 | Rabbit | 1:150 | Anti-Rabbit AlexaFluor 546 |
|  | Hoechst | N/A | 1:1000 | N/A |
|  | - | - | - | - |
|  | - | - | - | - |

**Supplementary Table S4.** 4i staining for KPCY used for spatial proximity analysis.

| Round | Antibody/Stain | Species | Dilution | Secondary |
| --- | --- | --- | --- | --- |
| **1** | Hypoxypore | Rabbit | 1:100 | Anti-Rabbit AlexaFluor 546 |
|  | GFP | Goat | 1:500 | Anti-Goat AlexaFluor 647 |
|  | Hoechst | N/A | 1:1000 | N/A |
| **2** | E-cadherin | Rat | 1:250 | Anti-Rat AlexaFluor 488 |
|  | Vimentin | Rabbit | 1:200 | Anti-Rabbit AlexaFluor 546 |
|  | Podoplanin | Syrian Hamster | 1:200 | Anti-Hamster AlexaFluor 647 |
|  | Hoechst | N/A | 1:1000 |  |

**Supplementary Table S5.** 4i staining for subcutaneous 0082T-HFM+HPAF-II tumors.

| Round | Antibody/Stain | Species | Dilution | Secondary antibody |
| --- | --- | --- | --- | --- |
| **1** | Vimentin | Mouse | 1:200 | Anti-Mouse AlexaFluor 546 |
|  | Hypoxyprobe | Rabbit | 1:100 | Anti-Rabbit AlexaFluor 488 |
|  | GFP | Goat | 1:100 | Anti-Goat AlexaFluor 647 |
|  | Hoechst | N/A | 1:1000 | N/A |
| **2** | RFP | Rabbit | 1:200 | Anti-Rabbit AlexaFluor 647 |
|  | Hoechst | N/A | 1:1000 | N/A |
|  | - | - | - | - |
|  | - | - | - | - |
| **3** | Podoplanin | Syrian Hamster | 1:200 | Anti-Hamster AlexaFluor 488 |
|  | COXIV | Rabbit | 1:500 | Anti-Rabbit AlexaFluor 546 |
|  | Hoechst | N/A | 1:1000 | N/A |
|  | - | - | - | - |

**Supplementary Table S6.**
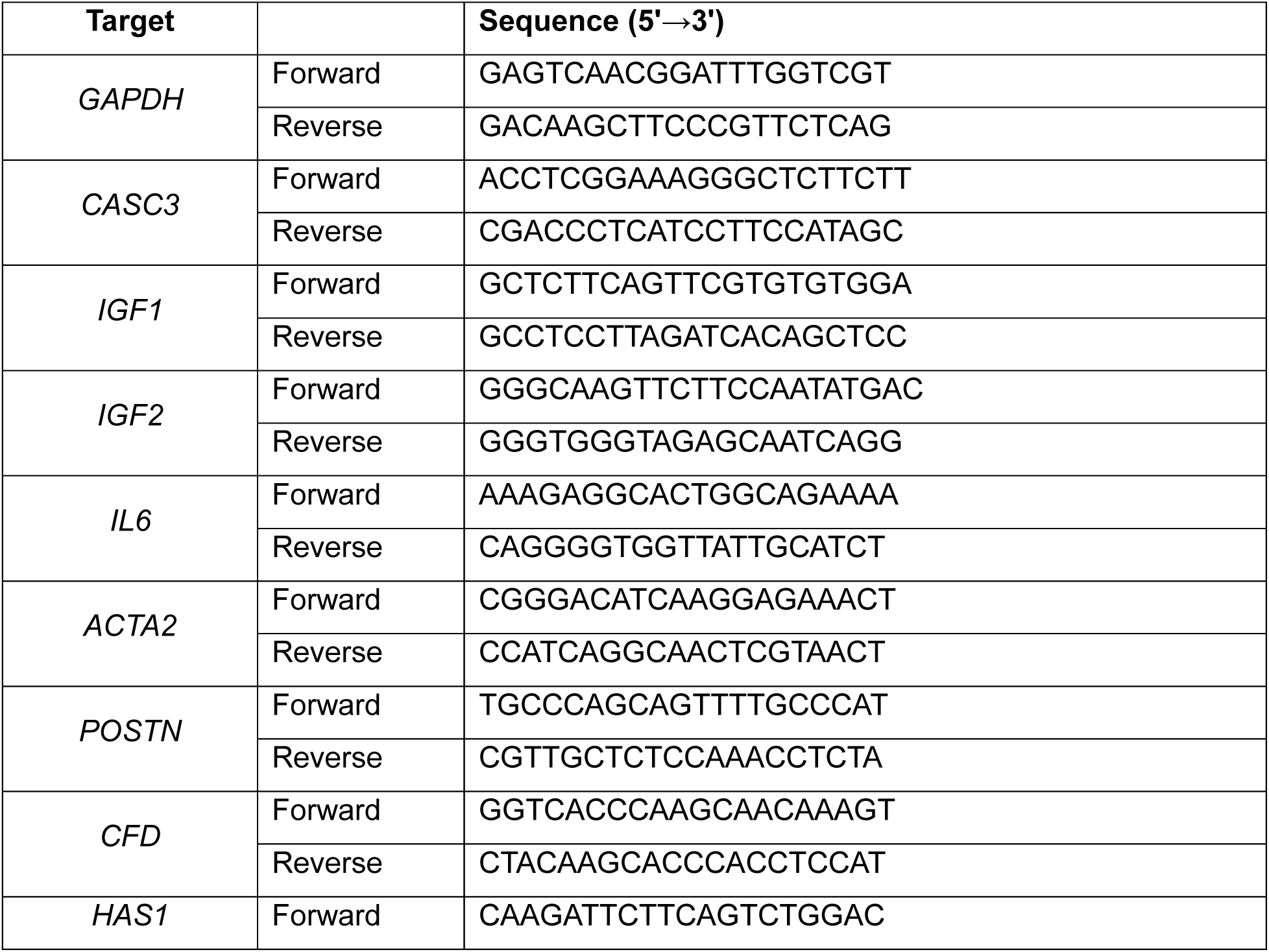
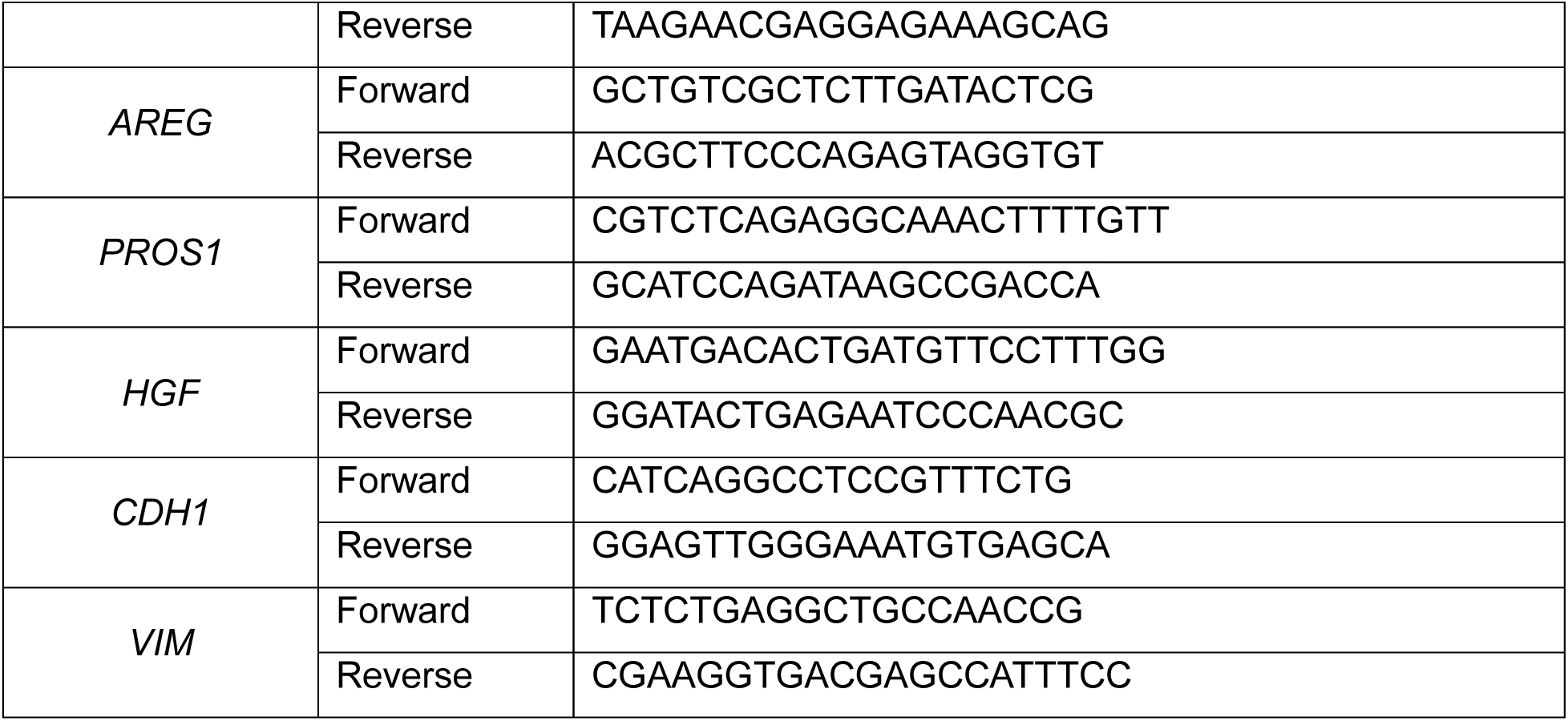
qRT-PCR primers.

**Supplementary Table S7.**
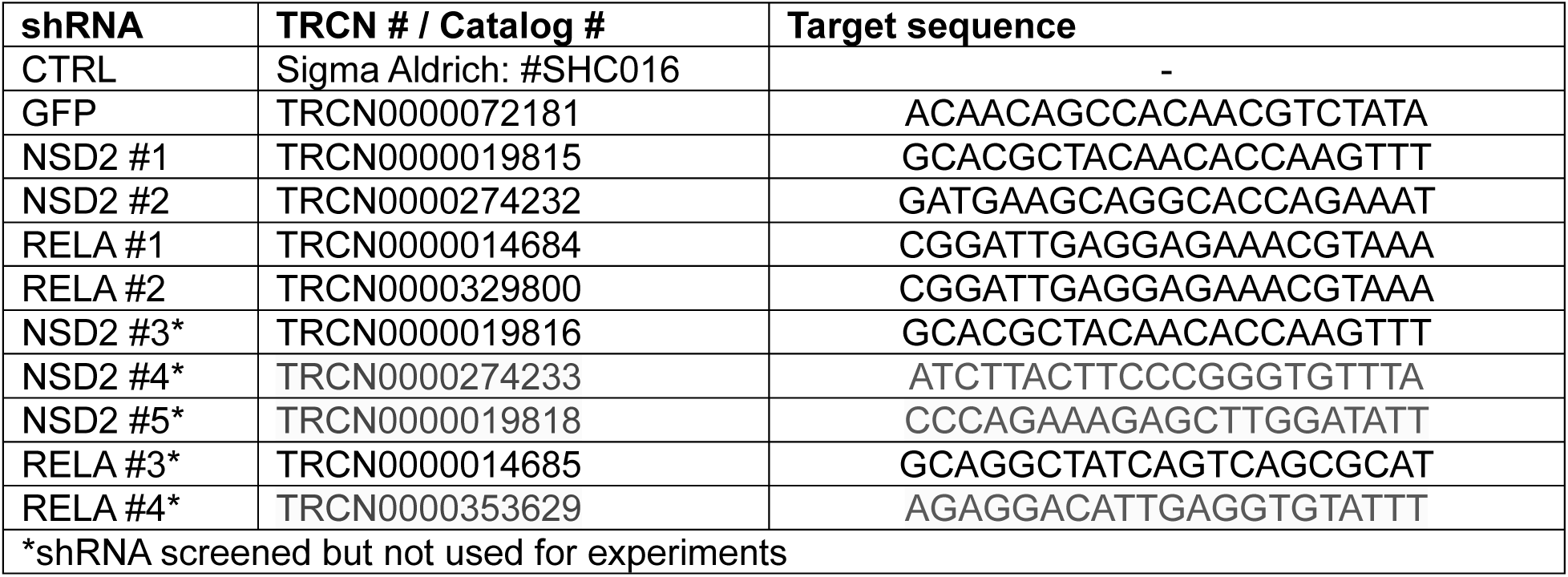
shRNA sequences.

